# Long-range communication between transmembrane- and nucleotide-binding domains does not depend on drug binding to mutant P-glycoprotein

**DOI:** 10.1101/2022.06.30.498271

**Authors:** Cátia A. Bonito, Ricardo J. Ferreira, Maria-José.U. Ferreira, Jean-Pierre Gillet, M. Natália D. S. Cordeiro, Daniel J. V. A. dos Santos

## Abstract

The modulation of drug efflux by P-glycoprotein (P-gp, ABCB1) represents one of the most promising approaches to overcome multidrug resistance (MDR) in cancer cells, however the mechanisms of drug specificity and signal-transmission are still poorly understood, hampering the development of more selective and efficient P-gp modulators. In this study, the impact of four P-gp mutations (G185V, G830V, F978A and ΔF335) on drug-binding and efflux-related signal-transmission mechanism was comprehensively evaluated in the presence of ligands within the drug-binding pocket (DBP), which are experimentally related with changes in their drug efflux profiles. The severe repacking of the transmembrane helices (TMH), induced by mutations and exacerbated by the presence of ligands, indicates that P-gp is sensitive to perturbations in the transmembrane region. Alterations on drug-binding were also observed as a consequence of the TMH repacking, but were not always correlated with alterations on ligands binding mode and/or binding affinity. Finally, and although all P-gp variants *holo* systems showed considerable changes in the intracellular coupling helices/nucleotide-binding domain (ICH-NBD) interactions, they seem to be primarily induced by the mutation itself rather than by the presence of ligands within the DBP. The data further suggest that the changes in drug efflux experimentally reported are mostly related with changes on drug specificity rather than effects on signal-transmission mechanism. We also hypothesize that an increase in the drug-binding affinity may also be related with the decreased drug efflux, while minor changes in binding affinities are possibly related with the increased drug efflux observed in transfected cells.

## 1. INTRODUCTION

Over-expression of membrane efflux pumps is intimately related to multidrug-resistance (MDR) phenomenom in cancer cells [1]. Among them, P-glycoprotein (P-gp, *ABCB1*) is the most studied so far, and it is capable to extrude a wide range of neutral and charged hydrophobic compounds) [1], through an ATP-dependent mechanism [2].

P-glycoprotein architecture comprises two transmembrane domains (TMDs), each one formed by six transmembrane α-helices (TMHs) packed in a pseudo 2-fold symmetry, and two cytoplasmic nucleotide-binding domains (NBDs), with both P-gp halves connected through a small peptide sequence (the “linker”; residues 627-688) [3]. The transmembrane and cytoplasmic domains of the P-gp efflux pump are physically linked to the respective NBD by coils bridging TMH6/NBD1 and TMH12/NBD2, and by short intracellular coupling helices (ICHs), located between TMHs 2/3 (ICH1-NBD1), 4/5 (ICH2-NBD2), 8/9 (ICH3-NBD2) and 10/11 (ICH4-NBD1), the latter involved in the TMD-NBD communication through non-covalent interactions [4,5]. Additionally, some studies indicated that ICHs may also play important roles in P-gp folding and maturation [6,7]. Oppositely, the drug-binding pocket (DBP) is located within the TMHs of both N- and C-terminal P-gp halves, and contains, at least, three distinct drug-binding sites (DBS): the modulator site (M-site) located at the top of DBP, and the substrate-binding sites (SBSs) H and R, named after Hoechst33342 and Rhodamine-123 respectively, located in the opposite side next to the inner leaflet of the lipid bilayer [8].

Although decades of research focus on the molecular basis of drug promiscuity and efflux, the mechanisms of drug specificity and efflux-related signal-transmission are still unclear, thus hampering the development of novel, more potent and selective P-gp modulators able to overcome MDR. Herein, one interesting approach is the study of P-gp variants experimentally related to altered drug-resistance phenotypes and/or changes in the basal and drug-stimulated ATPase activity. Our previous work [5], reported the impact of four hP-gp variants involved in MDR – G185V [9–14], G830V [14,15], F978A [16,17] and ΔF335 [18–21] – on the P-gp structure in the absence of ligands within the DBP (*apo* systems). In this paper, in order to understand if the P-gp conformational changes induce by these P-gp mutations affect drug-binding and efflux-related signal-transmission mechanism, the structural impact of these P-gp mutations was assessed in the presence of molecules, experimentally described as having alterations in their efflux profiles in these variants, bound to the reported DBSs (*holo* systems).

Similar to our previous work, as the glycine mutations G185V (TMH3) and G830V (TMH9) are located at the SBSs H and R, respectively, both variants will be named SBS-variants. On the other hand, the F978A (TMH12) and ΔF335 (TMH6) mutations, both lying at the M-site, will be referred as M-site variants. Following, the ligands were chosen considering: i) the location of the mutation at the TMDs, ii) the experimental and/or computational evidences for its putative DBS [8, 22–27], and iii) opposite changes in the drug efflux profiles by the same P-gp mutation [5]. Thus, colchicine (COL) and vinblastine (VIN), both predicted to bind at the H-site, were the substrates chosen for the G185V (H-site). Likewise, doxorubicin (DOX) and actinomycin D (ACT), both predicted to interact with the R-site, were the substrates selected for the G830V (R-site). On the other hand, since the mutations at the M-site (F978A and ΔF335) seem to strongly affect the SBSs properties and drug-binding [5], the structural effects of these mutations were analyzed in the presence of ligands in all three sites. Therefore, the P-gp modulator cyclosporine A (CYC, M-site), and the substrates ACT (R-site) and VIN (H-site) were the molecules selected for the F978A mutation. However, to the best of our knowledge, there is no information about a P-gp substrate that binds at the H-site and has altered drug-resistance profile in transfected cells harboring the ΔF335 mutation. That said, DOX (R-site) and two P-gp modulators, CYC and valspodar (VLS), both predicted to bind at the M-site, were the ligands chosen for this P-gp variant.

## 2. MATERIAL AND METHODS

### 2.1. Initial structures

From our previous study, five MD *apo* systems were used as template to build the MD *holo* systems, the refined and validated hP-gp wild-type (WT) and four hP-gp variants models (G185V, G830V, F978A and ΔF335). In each one the transporter is embedded in a lipid bilayer with 469 molecules of 1-palmitoyl-2-oleoylphosphatidylcholine (POPC), for which the lipid parameterization by Poger et al. [28,29] as previously described [5].

From our docking studies previously published [5], the top-ranked docking poses for the chosen ligands were selected and used to assemble the WT and variants MD *holo* systems. The molecules were parameterized in the PRODRG [30] online server and manually curated, according to the GROMOS96 force field [31–34], using AM1-BCC partial charges calculated with Antechamber 1.27 [35,36]. The protein-ligand-membrane systems were then solvated with 63.810 ± 31 single-point charge (SPC) water molecules and neutralized with 11 chlorine ions, for a total of 16 MD *holo* systems. These MD systems were the starting point for several short MD runs.

### 2.2. Free-energy calculations and analysis

After a short energy minimization run to minimize clashes between the ligand and the protein, a total of five replicates of 20-ns MD runs were performed for each system (N = 16), in a total of 1.6 μs of simulation time (0.1 μs each system). For each replicate initial velocities were assigned from a Maxwell-Boltzmann distribution at 303 K. To enhance sampling, for the last two replicates distinct snapshots were retrieved from the first MD run (at 18 ns and 19 ns, respectively). Only the last 10 ns of simulation time were considered as production runs and used for further analysis. Relative free-energies of binding (Δ*G*_bind_) were calculated using the *g_mmpbsa* tool [37], with an implicit membrane correction for polar solvation energies [38]. As in a previous paper, *gmx bundle*, *gmx hbond* [39] and *g_contacts* [40] tools were used to further evaluate changes in the transmembrane (TM) helical bundle, hydrogen bond (HB) networks, and protein interactions, respectively.

### 2.3. MD simulation parameters

All MD simulations were done with GROMACS v2016.x package [41]. All *NVT* equilibration runs were performed at 303 K using the Velocity-rescale (V-rescale) [42] thermostat. The Nosé-Hoover [43,44] thermostat and the Parrinello-Rahman [45] barostat for temperature (303 K) and pressure (1 bar), respectively, were applied in all *NpT* runs. Due to the presence of membrane, pressure equilibration was achieved through a semi-isotropic pressure coupling, with the systems’ compressibility set to 4.5 × 10^-5^ bar^-1^. All bond lengths were constrained using the LINCS [46,47] or SETTLE [48] (for water molecules) algorithms. The Particle Mesh Ewald (PME) with cubic interpolation [49,50] was employed with a cut-off radius of 12 Å for both electrostatic and van der Waals interactions, and an FFT grid spacing of 0.16 for long range electrostatics. Group-based and Verlet [51] cut-off schemes were applied for the calculation of non-bonded interactions on CPU or GPU, respectively.

## 3. RESULTS AND DISCUSSION

The analysis that will be presented and discussed in the following subsections relates to the behavior of the human P-gp WT and variants studied containing different allocrites inside the drug-binding sites.

### 3.1. Structural analysis of transmembrane domains of the hP-gp variants

To give additional insights on drug specificity and efflux-related signal-transmission mechanism, the structural impact of four selected P-gp mutations (G185V, G830V, F978A and ΔF335), experimentally related to changes in drug efflux and substrate specificity, were analyzed in the presence of molecules bound to each site inside the DBP (*holo* systems). The molecules chosen are described in literature as having altered efflux profiles in these P-gp variants—G185V (COL, VIN), G830V (ACT, DOX), F978A (ACT, VIN, CYC), and ΔF335 (CYC, VLS, DOX) (Table 1).

**Table 1.** Structural impact of mutations in the TMH repacking for the *holo* hP-gp variants.^a^

| Mutation-DBS | Molecule-DBS | Repacking effects |
| --- | --- | --- |
| **G185V-H** | COL-H | +++ |
|  | VIN-H | ++ |
| **G830V-R** | DOX-R | +++ |
|  | ACT-R | +++ |
| **F978A-M** | CYC-M | +++ |
|  | ACT-R | +++ |
|  | VIN-H | + |
| **ΔF335-M** | CYC-M | + |
|  | VLS-M | + |
|  | DOX-R | + |

<sup>a</sup> Positive variation – lowest impact (+); highest impact (+++). The P-gp variants *holo* systems were compared both with the WT and between them.

Mutations [5] and/or the presence of ligands within the DBP may directly affect the TMDs architecture [52]. Thus, the rearrangement of the transmembrane helices (TMHs) was assessed using the *gmx bundle* for the Pgp variants *holo* systems and compared to the WT (*holo* system). All the significant changes in the bundle parameters are summarized in the Supporting Information (SI) (*Tables S1-S24 and Figures S1-S48*). To organize and simplify the results described below, the studied P-gp systems will be encoded and referred as “variant-DBS1_ligand-DBS2”, where DBS1 is DBS where the mutation is located and DBS2 the DBS where the molecule is reported to bind.

Overall, the bundle parameters calculated for all P-gp variants were found to have distinct shifts from the WT, when in the presence of the respective ligands. Most remarkably, the changes were larger when the helices that form the DBP portals (4/6 and 10/12) and the “crossing helices” (4/5 and 10/11), directly connecting the TMD1 to NBD2 and TMD2 to NBD1 respectively [53], are involved. However, distinguishable structural effects in the TMH rearrangement were still observed between P-gp variants. The SBS-variants G185V and G830V, in the presence of the respective ligands, revealed deeper changes in the TMH repacking affecting both P-gp halves. Firstly, COL in the G185V-H_COL-H system induced a stronger impact in the TMDs architecture than VIN in the G185V-H_VIN-H system. Similarly, the F978A variant showed a striking TMH repacking, more severe in the F978A-M_CYC-M and F978A-M_ACT-R systems. On the other hand, the total repacking observed in the ΔF335 variant seemed to be similar in all *holo* ΔF335 systems.

The structural impact of a specific ligand in two different P-gp variants was determined (Table 1). Herein, a stronger effect in the TMH repacking was observed in the G185V-H_VIN-H and G830V-R_DOX-R systems than those observed for the M-site variants in the F978A-M_VIN-H and ΔF335-M_DOX-R systems, respectively. On the other hand, ACT induced a similar impact in the TMH rearrangement in both G830V-R_ACT-R and F978A-M_ACT-R systems. Moreover, CYC seemed to have larger shifts in the bundle parameters in the F978A-M_CYC-M system than for the ΔF335-M_CYC-M system. Finally, when compared to the P-gp variants *apo* systems [5], more severe changes in the TMDs architecture were observed for all *holo* P-gp variants. Nonetheless, the presence of ligands in the M-site variants seemed to have a larger effect in the TMH repacking than those reported for the respective *apo* systems.

Altogether, the analysis of the helical bundle indicates that the studied P-gp mutations, when in the presence of ligands within the DBP, i) severely affect the native spatial position of the helices that form the DBP portals, which may alter the access of drugs to the internal cavity or their release to the membrane [1], and ii) induces changes in the helical bundle concerning the “crossing helices”, crucial for the NBD dimerization process upon ATP-binding [53]. Additionally, the presence of molecules seems to induce a more severe impact in the TMHs rearrangement than those observed in the respective *apo* systems [5], indicating a TMDs adjustment and response to the presence of ligands.

### 3.2. Characterization of protein-ligand interactions

Since all the above mutations changed the TMDs architecture in variable degrees, their impact on the ligand-binding affinity was further evaluated. Relative free-energies of binding (Δ*G*_bind_) were estimated using the *g_mmpbsa* (only available in GROMACS v5.x) tool and compared to the WT *holo* systems (*Supporting Information, Figures S49-S52*).

Although a severe TMH repacking was observed in all P-gp variants, the binding affinity was affected differently (Table 2). While no significant changes were found in the G185V-H_COL-H system, a substantial increase in binding affinity was observed in both G185V-H VIN-H and F978A-M_VIN-H systems. Oppositely, no considerable effects were observed in the calculated Δ*G*_bind_ values for the G830V-R DOX-R and ΔF335-M_DOX-R systems. No significant changes in binding affinities were also found for the G830V-R_ACT-R system. On the other hand, although observing a slight increase in the ACT-binding affinity in the F978A-M_ACT-R system, no clear conclusions could be made about the possible changes in the molecule’s affinity. Finally, an increase on the ligand-binding affinities were observed in the F978A-M_CYC_M, ΔF335-M CYC-M and ΔF335-M_VLS-M systems.

**Table 2.** Structural impact of mutations in the ligand-binding affinity for the P-gp variants *holo* systems.

| Mutation-DBS | Molecule-DBS | Binding affinity impact<sup>a</sup> |
| --- | --- | --- |
| **G185V-H** | COL-H | NS |
|  | VIN-H | ↑↑↑ |
| **G830V-R** | DOX-R | NS |
|  | ACT-R | NS |
| **F978A-M** | CYC-M | ↑↑ |
|  | ACT-R | ↑ |
|  | VIN-H | ↑↑↑ |
| **ΔF335-M** | CYC-M | ↑↑ |
|  | VLS-M | ↑↑ |
|  | DOX-R | NS |

<sup>a</sup> Positive variation – lowest impact (+); highest impact (+++); not significant (NS). The P-gp variants *holo* systems were compared both with the WT and between them.

To understand how the severe TMH rearrangement observed in all P-gp variants affected ligand-binding affinity differently, specific protein-ligand interactions were additionally assessed using the *g_contacts* (from GROMACS v4.6.7) and *gmx hbond* tools, and again compared to the WT protein *holo* systems. Mean contact frequencies ≥ 0.5 and variations above 10% were considered as significant and are depicted in SI (*Supporting Information, Tables S25-S27*).

According to our previous studies, the TMH repacking induced by these P-gp mutations led to severe changes in both structure and residues distribution at the DBSs, thus having a strong impact on size and polarity [5]. Herein, and concerning the G185V-H_COL-H system, colchicine, initially located at the bottom of the H-site, shifted from its original binding position in WT, losing interactions with TMHs 10/12 while establishing new ones with TMHs 6/11 (Figure 1).

**Figure 1.**
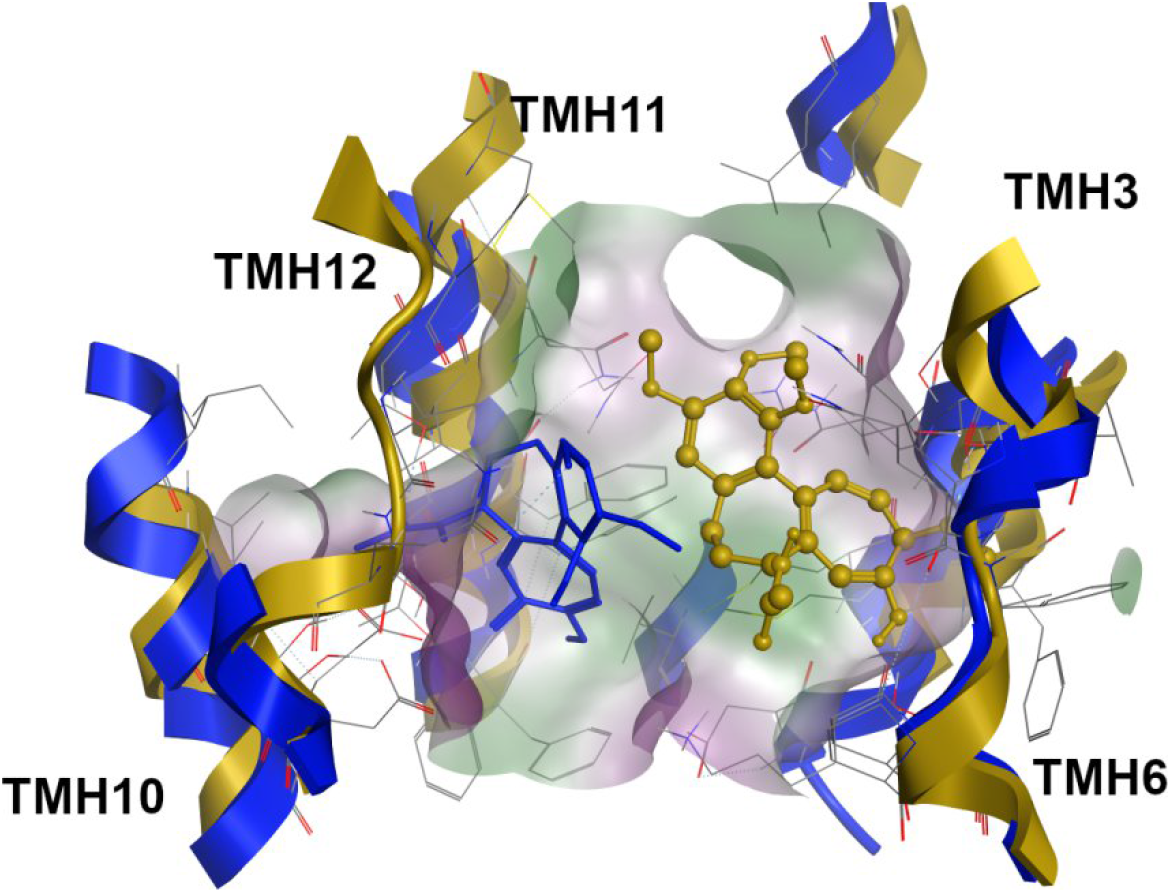
Superimposition of the colchicine-bound systems at the H-site for WT (blue, licorice) and G185V variant (dark yellow, ball-and-stick).

However, and according to the ratio between the number of significant protein-ligand contacts and the average of the total number of contacts (hereby named contact efficiency ratio – CE, *Supporting Information, Table S28*), similar values were observed in both WT-H_COL-H and G185V-H_COL-H systems, thus indicating that although COL established new contact points with residues from other TMHs, no significant impact in binding affinity was observed for this ligand. In the same way, a similar CE ratio was observed between the WT-R_DOX-R and the variant *holo* systems G830V-R_DOX-R and ΔF335-M_DOX-R.

Regarding doxorubicin, a common binding mode between the WT and both G830V and ΔF335 variants was observed, in which the tetracyclic core is found in close contact with hydrophobic surface patches – by interacting with residues from TMHs 7 and 9 – in the respective pockets (Figure 2). These results indicate that the binding mode for DOX is not substantially affected by these mutations, and no significant alterations in the DOX binding affinity were observed for these variants. Oppositely, a great increase in the vinblastine contact efficiency ratio was observed in both G185V-H_VIN-H and F978A-M_VIN-H systems. The visual inspection showed that VIN was deeply buried in the H-site in both P-gp variants, which may account for stronger interactions and, concomitantly, to the higher binding affinity observed for this molecule.

**Figure 2.**
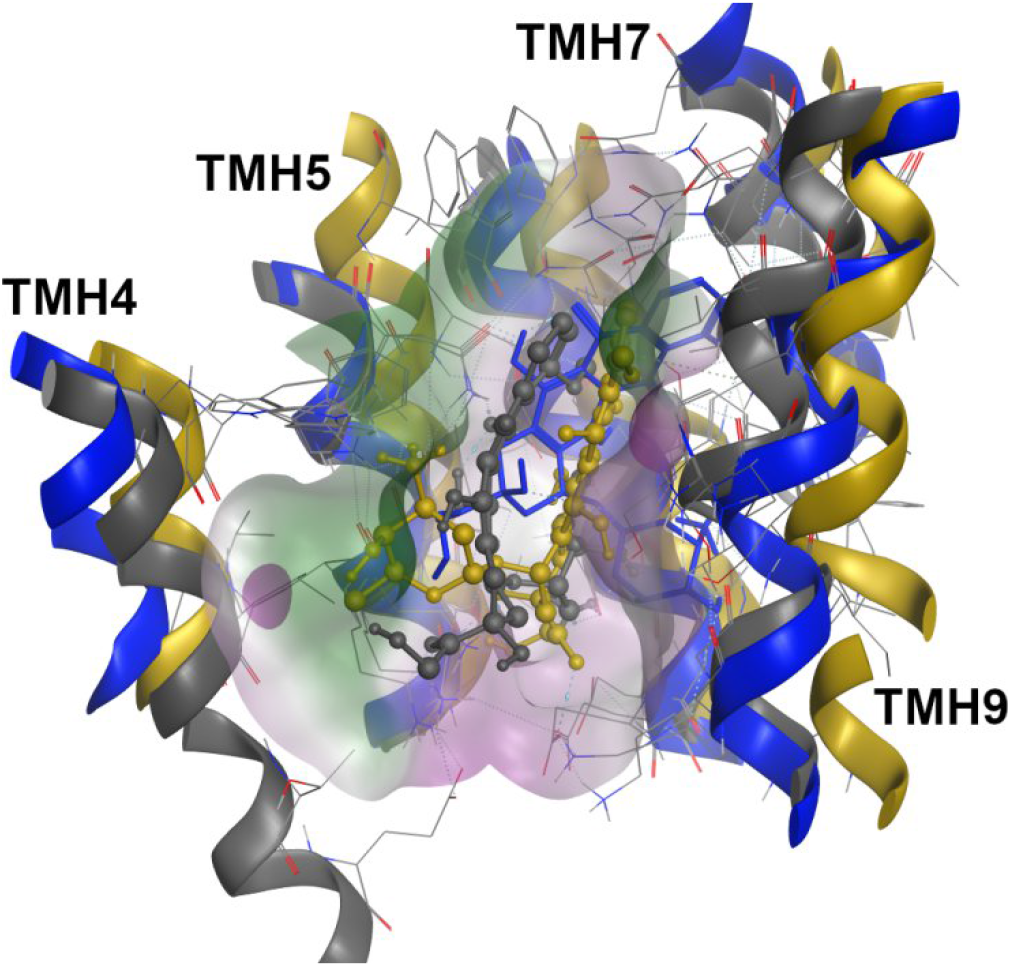
Superimposition of the doxorubicin-bound systems at the R-site for the WT (blue, licorice), G830V (dark yellow, ball-and-stick) and ΔF335 (gray, ball-and-stick) variants.

Quite interestingly, a complete distinct binding mode was observed in the ACT in the G830V-R_ACT-R system. The visual inspection revealed severe alterations in the ACT-binding mode, with a large region of the molecule protruding from the R-site towards the middle of the DBP and, thus exposed to the surrounding environment (Figure 3).

**Figure 3.**
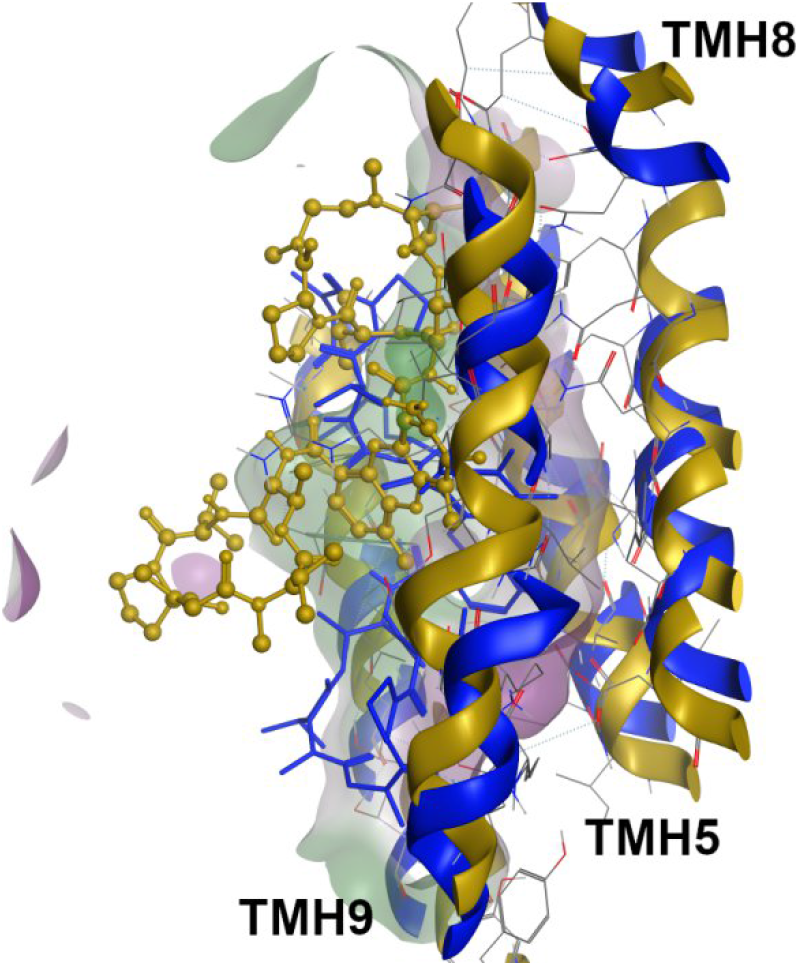
Superimposition of the actinomycin-bound systems at the R-site for the WT (blue, licorice) and G830V variant (dark yellow, ball and stick).

Since the contact efficiency (CE) ratio was higher in the G830V-R_ACT-R system than in WT-R_ACT-R but without significant change in binding energy, the loss of contacts between a large region of the molecule and the protein may explain the minor impact in the ACT Δ*G*_bind_ values observed for this variant i.e the molecule changes the interaction pattern loosing interactions by making fewer but stronger ones. Oppositely, a visual inspection of the F978A variant showed that ACT remained deeply buried in the R-site as observed in WT, thus showing a similar contact efficiency ratio and, as expected, no changes in the ACT binding affinity.

Lastly, an increase in the CE ratio was observed for both F978A-M_CYC-M and ΔF335-M_CYC-M systems. Again, visual inspection showed that the structural changes at the M-site in the F978A variant prompted CYC to change the conformation of its macrocycle, shifting one of the isoleucine side-chains into the more hydrophobic environment at the center of the macrocyclic core (Figure 4) and therefore improving the CYC binding affinity.

**Figure 4.**
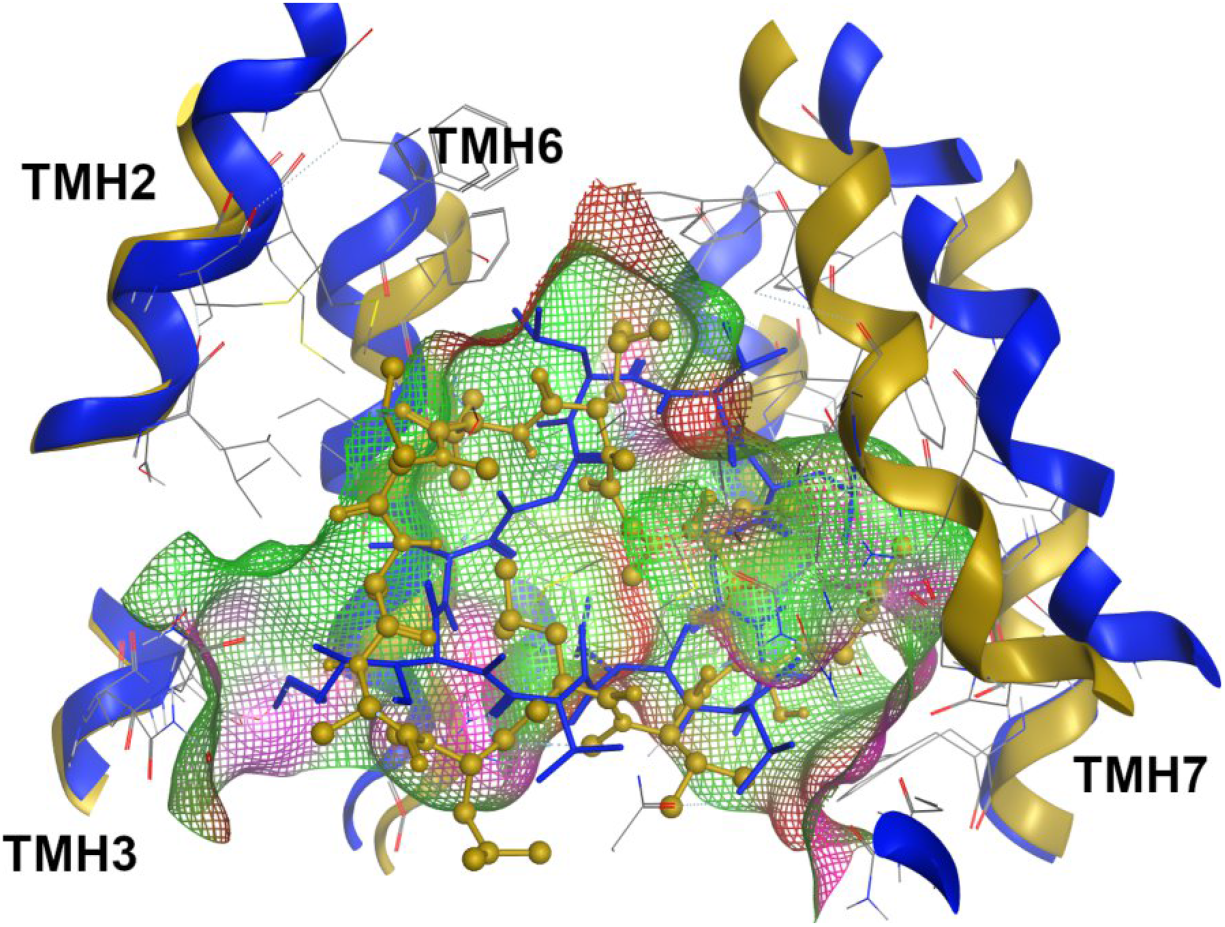
Superimposition of the cyclosporine-bound systems at the M-site for the WT (blue, licorice) and F978A variant (dark yellow, ball-and-stick).

Regarding the ΔF335-M_CYC-M system, no considerable changes in the CYC binding mode was observed in respect to WT, with all hydrophobic groups being protected from the solvent. However, the increase in the CYC binding affinity observed for these variants allow to infer that small changes of conformation in a large ligand may play important roles in binding affinity. Likewise, valspodar in the ΔF335-M_VLS-M system was also found to be deeply buried inside the M-site, which favors the reinforcement of the VLS-ΔF335 interactions, and consequently, its binding affinity. These results could be explained by the lower impact in the M-site volume and polarity reported for the F978A and ΔF335 variants *apo* systems, when compared to the SBSs where the structural impact of these mutations was higher [5].

Altogether, the above data indicate that these P-gp mutations seem to have an impact on the binding energy, ligand-binding modes and/or on the protein-ligand interactions. Moreover, although the mutations have an undeniably distinct but strong structural impact in the TMH repacking, similar effects on the binding energy observed for the same molecule across two different mutations (VIN-H in G185V and F978A; ACT-R in G830V and F978A; DOX-R in G830V and ΔF335; CYC-M in F978A and ΔF335), suggest that the molecular properties of these ligands may play important contributions to their binding affinity rather than the severity of the TMH repacking induced by these mutations. This remark seems to be independent of the DBS where the molecule binds.

### 3.3. Interactions between coupling helices and nucleotide-binding domains

All the herein considered Pgp variants are experimentally described as having altered drug efflux profiles and changes in the basal and/or drug-stimulated ATPase activity [9–21]. Therefore, as the ICH-NBD residue interactions are increasingly considered to be key players in signal-transmission and efflux-related conformational changes [7,54–57], a thorough evaluation of the ICH-NBD contacts was performed and compared to the WT *holo* systems. The total number of contacts was estimated by the *gmx hbond* module (*Supporting Information, Figures S53-S60*).

Overall, all P-gp *holo* systems revealed changes in the mean total number of contacts at the ICH-NBD signal-transmission interfaces. However, while in most cases a reinforcement in the total number of contacts was observed at the ICHs-NBD1 interfaces, a negative impact in the total number of contacts was observed in the ICHs interacting with NBD2 (Table 3).

**Table 3.** Structural impact of mutations in the total number of contacts at the ICHs-NBD interfaces for the P-gp variants *holo* systems.

| Molecule-DBS | Mutation-DBS | ICHs-NBD1 |  | ICHs-NBD2 |  |
| --- | --- | --- | --- | --- | --- |
|  |  | ICH1 | ICH4 | ICH2 | ICH3 |
| **G185V-H** | COL-H | ↑↑<sup>a</sup> | NS | NS | ↓<sup>b</sup> |
|  | VIN-H | NS | ↑ | NS | ↓ |
| **G830V-R** | ACT-R | ↑↑ | NS | ↓ | ↓ |
|  | DOX-R | ↑ | ↑ | ↓ | ↓ |
| **F978A-M** | VIN-H | ↑↑ | ↑↑ | NS | NS |
|  | ACT-R | ↑ | ↑ | ↓↓ | NS |
|  | CYC-M | ↑ | ↑↑ | NS | NS |
| **ΔF335-M** | DOX-R | ↑↑ | ↑↑ | ↓↓ | NS |
|  | CYC-M | ↑↑↑ | NS | ↓ | ↓ |
|  | VLS-M | ↑↑ | ↑ | NS | NS |

<sup>a</sup> Positive variation — the lowest impact (↑), the highest impact (↑↑↑); <sup>b</sup> negative variation (↓), — the lowest impact (↓), the highest impact (↓↓); NS — not significant. The P-gp variants *holo* systems were compared both with the WT and between them.

As all P-gp mutations in the presence of ligands seemed to affect the total number of contacts at the ICH-NBD interfaces, we aimed for the identification of which residue pairs were mostly involved through the evaluation of their mean contact frequencies, using the *g_contacts* tool. Only mean contact frequencies ≥ 0.5 and variations above 10% were considered meaningful (*Supporting Information, Tables S28-S32)*. For clarity purposes, the analysis of each ICH-NBD interface will be described separately:

#### 3.3.1. ICH-NBD1 interfaces

Starting from the N-terminal P-gp halve, the first ICH-NBD interface is the ICH1-NBD1. Herein, all P-gp variants *holo* systems showed a significant increase in the mean contact frequencies between I160 and the NBD1 residue L443, described as important for ATP-binding [57–59]. On the contrary, a negative impact in the residue interactions was observed in most of the P-gp variants *holo* systems between D164-K405, also involved in the TMD-NBD communication pathway [52]. When concerning the hydrogen bond network, a negative effect in the HB lifetime was also observed in both G830V-R_ACT-R and F978A-M_ACT-R systems as well as in all ΔF335 variant *holo* systems. A decrease in the HB lifetime was also inferred from the G185V-H_VIN-H, but when in the presence of colchicine a significant reinforcement of the HB network in the G185V-H_COL-H system was observed.

On the other hand, all the P-gp variants showed clear differences in the residue interactions at the ICH4-NBD1, the second signal-transmission interface involving NBD1. Regarding the SBS-variants, similar effects in the residue interactions were observed in the G185V *holo* systems, mainly reinforcing the contact frequencies between Q912-R467, with R467 being involved in the propagation of the conformational changes [60]. Oppositely, a strong negative impact in the mean contact frequencies was observed for the residue pair T911-R467. A negative variation in the contact frequencies was also verified to occur for the residue pairs S909-Q441 and Q912-R464, with S909 being involved in the activation and ATPase stimulation when in the presence of drugs and/or lipids [57, 61]. In the G830V-R_ACT-R system, the presence of ACT again induced a strong negative effect in the contacts involving the R905-Q438, T911-R467, and Q912-R464 residue pairs. Quite interestingly, mutational studies already had suggested that R905 is a pivotal residue in the activation and ATPase stimulation when in the presence of drugs and/or lipids [57, 61]. Surprisingly, an opposite trend was found for both M-site variants. Both the F978A and ΔF335 *holo* systems showed a significant reinforcement in the mean contact frequencies between the R905-Y401/Q438 and V908/T911-R467 residue pairs, with Y401 (NBD1) again being important for ATP binding [53]. Regarding hydrogen bonds, the G830V-R_ACT-R system was the only one that showed a dramatic increase in the HB lifetime while a negative impact in the calculated HB parameters was observed in all other P-gp variants *holo* systems.

#### 3.3.2. ICH-NBD2 interfaces

At the C-terminal P-gp halve, the first signal-transmission interface is the ICH2-NBD2. Herein, all P-gp variants *holo* systems showed a decrease in the mean contact frequencies between the residues A266 (ICH2) and F1086 (NBD2), a critical residue pair for coupling of ATP-binding to conformational changes in the TMDs [6,7]. Nonetheless, substantial differences in residue pair interactions were observed between the P-gp variants. For both G185V-H_COL-H and G185V-H_VIN-H systems, the presence of these ligands affected the R262-F1086 (positive variation) and I265-R1110 (negative variation) residue pairs, being R262 (ICH2) an important residue in the propagation of the conformational changes [60]. However, a noteworthy reinforcement in the mean contact frequencies between T263-Q1118 and T263-D1200 residue pairs was observed only for both G185V *holo* systems. Accordingly, Q1118 is an important residue for ATP-binding/hydrolysis and Mg^2+^ binding since mutations in this residue, located within the Q-loop, blocked drug-stimulated ATPase activity [62,63]. In contrast, the presence of ACT and DOX in the G830V-R_ACT-R and G830V-R_DOX-R systems had an overall negative impact in the mean contact frequencies involving the R262-Q1081 and T263-S1117 residue pairs. Regarding the M-site variants, a negative variation in the mean contact frequencies was observed in the F978A variant *holo* systems, namely between R1110 (NBD2) and residues I265/F267 (ICH2), described as involved in P-gp maturation and activity [6,7,64]. Similarly, the presence of DOX in the ΔF335-M_DOX-R system negatively affected most of the identified ICH-NBD contact points.

Changes in the ICH-NBD hydrogen-bond network were additionally observed for the G185V and M-site variants *holo* systems. While the G185V-H_COL-H system showed a decreased in the HB lifetime, VIN on the other hand reinforced the calculated HB parameters in the G185V-H_VIN-H system. Opposite effects were observed between CYC and VLS in their respective P-gp variants: while the HB lifetime was negatively affected by the presence of CYC in both F978A-M_CYC-M and ΔF335-M_CYC-M systems, the presence of VLS in the ΔF335-M_VLS-M system instead reinforced the HB network.

Finally, most of the P-gp *holo* systems also showed negative effects in the residue interactions at the ICH3-NBD2, the second signal-transmission interface interacting with NBD2. Most particularly, a significant decrease in the mean contact frequencies was observed between the residue pairs V801/S802 and the NBD2 residue Y1087, which is thought to play an important role in P-gp activity and folding [56]. Moreover, the Y1087 residue was also predicted to be the key residue in the ICH3-NBD2 communication [55]. Similarly, and concerning the HB network, a general decrease in the HB lifetime was observed in the *holo* systems of the G185V and M-site variants.

Altogether the results indicate that the studied P-gp mutations *holo* systems i) have long-range effects in the residue interactions at the ICH-NBD signal-transmission interfaces, ii) induce asymmetries in the ICH-NBD residue interactions and iii) affect the mean contact frequencies of important residue pairs experimentally related to drug-stimulated ATPase activity, P-gp maturation and/or folding.

## MECHANISTIC INSIGHTS

P-glycoprotein over-expression is one of the most relevant multidrug-resistance mechanisms in cancer cells [65]. To develop compounds with high selectivity and efficacy towards P-gp, a deeper knowledge underlying the mechanisms of drug specificity and efflux-related signal-transmission is needed. In our previous work, a refined hP-gp WT model and four P-gp variants (G185V, G830V, F978A, and ΔF335), experimentally related to changes in substrate specificity, basal and drug-stimulated ATPase activity were built in the *apo* state inward-facing conformation, and their structural impact in the TMDs and ICH-NBD interactions was thoroughly evaluated [5]. In the present study, these P-gp models were used to assess the structural effects of these mutations on drug-binding and efflux-related signal-transmission mechanism in the presence of ligands within the DBP, that are also experimentally described as having altered efflux patterns.

Taken together, our data indicate that these P-gp mutations in the presence of ligands within the DBP induce structural effects in the TMDs by affecting i) the DBP portals 4/6 and 10/12 with the possibility for compromising the access and release of substrates, and ii) the “crossing helices” 4/5 and 10/11 able of compromising the NBD dimerization upon ATP-binding. Although these findings are in agreement with those reported in our P-gp *apo* systems [5], the impact on the TMH repacking seem to be stronger in the P-gp *holo* systems, suggesting that P-gp is sensitive to the presence of ligands. Additionally, our results are also in agreement with the experimental studies performed by Loo and Clarke [56]. Accordingly, substrates bind to P-gp through a “substrate-induced fit mechanism”, where the size and shape of the substrate induce rearrangements in the TMHs [56]. Furthermore, the rearrangement of the TMH residues’ side-chains due to repacking affect the protein-ligand interactions and/or binding affinity. On the other hand, the presence of mutations and/or ligands induces variations in the total number of contacts at the ICH-NBD interfaces, possibly affecting the TMD-NBD communication, and consequently altering drug efflux. Lastly, the impact in the mean contact frequencies of specific ICH-NBD residue pairs, experimentally described as involved in the signal-transmission mechanism, folding and ATP-binding [64], offers a possible explanation for the alterations in drug efflux reported for these P-gp variants.

Interestingly, most of the P-gp variants *holo* systems showed asymmetrical changes in the total number of contacts at the ICH-NBD interfaces, namely a reinforcement in the number of contacts at the ICHs-NBD1, while a decrease in the contacts was observed in the ICHs connecting NBD2. Two experimental studies recently demonstrated that i) the presence of cholesterol in the membrane induces asymmetries between NBDs, involving the ICH-NBD interfaces [66], and ii) there are common regions affected in a similar manner, but diverging in the post-hydrolysis state, especially in ICHs 3 and 4. Concerning the former, if taken into account that cholesterol is also a P-gp substrate [66], the latter corroborates our hypothesis that the presence of the selected ligands within the DBP exacerbate the asymmetries in the contacts at the ICH-NBD interfaces. The fact by which similar changes in the residue’s interactions at the ICH-NBD interfaces were also reported in these P-gp variants *apo* systems [5], especially concerning the M-site variants, additionally suggests that i) such alterations can be induced solely by the mutation, independently of the presence of molecules within the DBP, and ii) such alterations can be further exacerbated by the presence of molecules bound at the SBSs.

Nevertheless, important structural differences were observed between the P-gp variants *holo* systems. Regarding the G185V (TMH3) mutation, experimental data showed that it conferred increased resistance to COL in transfected cells, with an increase in the COL-stimulated ATPase activity, and a 3.6-fold decrease in COL binding. Oppositely, a decreased resistance to VIN was reported for this P-gp variant, with a 3.8- to 5.5-fold increase in the VIN binding [15]. Although no clear conclusions could be made about the possible changes in the COL binding affinity in the G185V-H_COL-H system, an indubitably increase in the VIN binding affinity was observed in the G185V-H_VIN-H system, in agreement with the experimental data.

Nonetheless, the presence of COL induced a reinforcement of the residue’s interactions and HB network at the ICH1-NBD1, something that was not observed when in the presence of VIN. As ICH1 establishes interactions with the A-loop residues of NBD1 and the adenine group of ATP [67], we hypothesized that the reinforcement of the ICH1-NBD1 interactions may be related to the increase in COL-stimulated ATPase activity reported in the experimental studies and, therefore, with the increased resistance to COL. Furthermore, a recent study emphasized the role of the conserved ICH1-NBD1 interface as critical for the cross-talk between the TMD-NBD domains in ABCG2, another relevant ABC transporter in the MDR in cancer cells [68]. On the other hand, the decrease in the VIN efflux experimentally observed in this P-gp variant, in our models, is solely due to the sharp increase in the VIN binding affinity. A previous experimental study performed by Loo and Clarke [17] focused on another single point mutation, F335A, which correlated the increase in the VIN binding affinity within the DBS with a decrease in the VIN efflux. According to the authors, the release of VIN during the transport cycle was postulated to be impaired due to its increased affinity at the DBS, and the herein published model corroborates those assumptions.

Located at the opposite P-gp halve, the G830V mutation (TMH9) in the presence of DOX (G830V-R_DOX-R) did not seem to significantly affect DOX binding affinity, as observed in the G185V-H_COL-H system. Although no more conclusions could be retrieved from our data, it clearly suggests that even minor effects in the ligands binding affinity may be related with the increased drug efflux. In contrast, and despite the overall increase in the contact efficiency ratio – which was expected to contribute for increasing the ACT-binding affinity –, the severe changes in ACT binding mode observed in the G830V-R_ACT-R system seemed to act as a “counter-weight”, having as consequence a non-significant change in the Δ*G*_bind_ values. Therefore, we suggested that in this specific case, the decrease in the ACT efflux is not related with the increase binding affinity as observed with VIN, but to the ACT unfavorable binding mode at the R-site, which may reduce the ability for the G830V variant in transport ACT. This hypothesis could be supported by experimental studies, where only a slight decrease resistance to ACT (0.29-fold) in respect to WT was observed in this P-gp variant [15].

Overall, it becomes clear that the binding of substrates at the H and R sites has similar effects on drug efflux in both glycine variants. Additionally, these findings also corroborate our previous studies, where we suggested that G185 and G830 residues, located at opposite halves have equivalent roles in P-gp function, likely more involved in drug binding [5]. Finally, and although no significant changes in the DOX-binding affinity in the ΔF335-M_DOX-R system, both F978A (TMH12) and ΔF335 (TMH6) mutations tend to induce an increase on ligand-binding affinities, which seems to be independent of the DBS where the molecules interact. Therefore, we hypothesize that for certain ligands, the increase in drug binding affinity within the DBS is a possible mechanism underlying the decreased drug efflux.

## AUTHOR CONTRIBUTIONS

C.A.B., R.J.F. and D.J.V.A.S. conceived the experiment(s), C.A.B. conducted the experiment(s), C.A.B., R.J.F., and D.J.V.A.S. analyzed the results and wrote the paper. C.A.B., R.J.F., D.J.V.A.S., M.J.U.F., J.-P.G. and M.N.D.S.C. reviewed the manuscript. All authors agreed with the final version of the manuscript.

