## Supplementary Information for "Long-range communication between transmembrane- and nucleotide-binding domains does not depend on drug binding to mutant P-glycoprotein"

### TABLE OF CONTENTS

**Tables S1-S24 and Figures S1-S48.** Results from the *gmx bundle* analysis.

**Figures S49-S52.** Characterization of protein-ligand interactions in the human P-gp variants (*g_mmpbsa*)

**Tables S25-S27**. Detailed results concerning protein-ligand interactions (*g_contacts* and *gmx hbond* tools)

**Table S28.** Protein-ligand contact efficiency ratio for the WT and P-gp variants.

**Figures S53-S60.** Total number of contacts at each ICH-NBD interface for the WT model and all P-gp *holo* systems and variation in the total number of contacts found at each ICH-NBD interface, for all P-gp variants, when compared to the WT.

**Tables S29-S32.** Variation in the contact frequencies and hydrogen bond (HB) parameters between the ICH-NBD residues for the WT and all variants (*holo* systems).

**Table S1.** Variation in the axis distance observed in the *holo* G185V variant transmembrane helices (compared to the *holo* WT; +, positive mean changes; –, negative mean changes).

| | TM1 | TM2 | TM3 | TM4 | TM5 | TM6 | TM7 | TM8 | TM9 | TM10 | TM11 | TM12 |
|---|---|---|---|---|---|---|---|---|---|---|---|---|
| G185V-H_COL-H | ns* | ns | + | + | - | ns | + | - | - | + | - | + |
| G185V-H_VIN-H | ns | ns | + | + | - | ns | + | - | - | + | - | + |

* ns, not significant

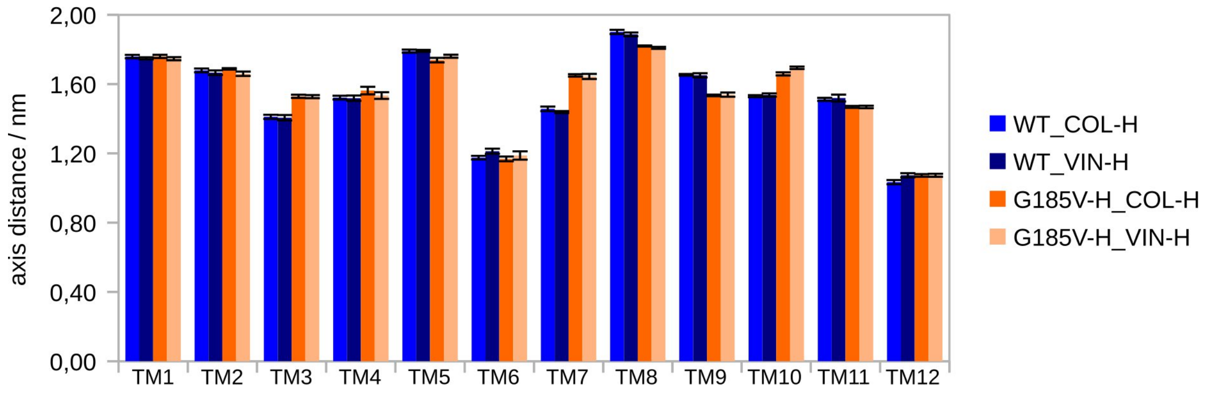


**Figure S1:** Comparison of individual transmembrane axis distance for the WT and G185V variant (*holo* systems).

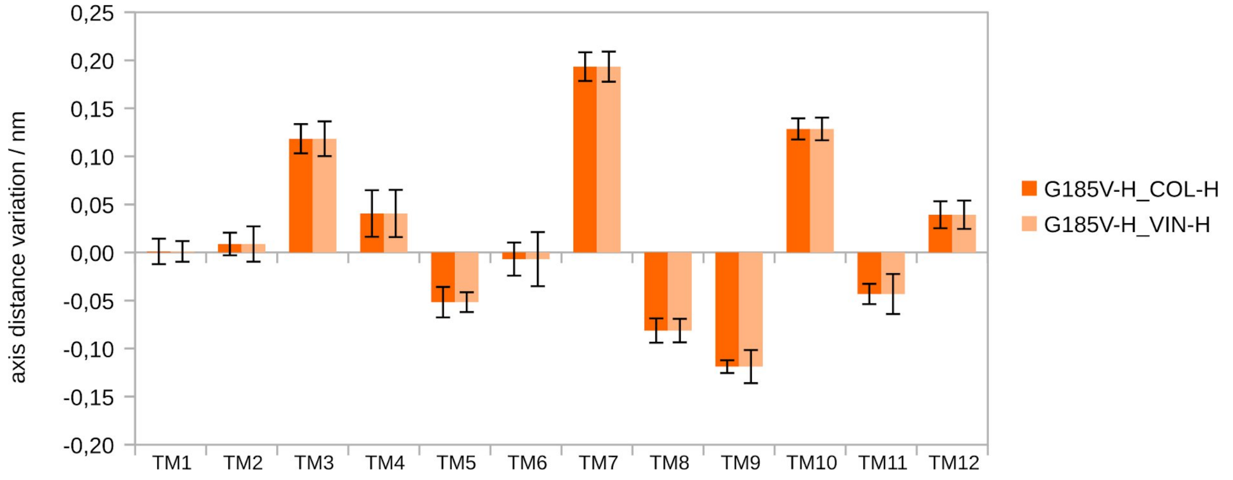


**Figure S2:** Variation in the transmembrane axis distance for the *holo* G185V variant (difference from the *holo* WT).

**Table S2.** Variation in the axis length observed in the *holo* G185V variant transmembrane helices (compared to the *holo* WT; +, positive mean changes; –, negative mean changes).

| | TM1 | TM2 | TM3 | TM4 | TM5 | TM6 | TM7 | TM8 | TM9 | TM10 | TM11 | TM12 |
|---|---|---|---|---|---|---|---|---|---|---|---|---|
| G185V-H_COL-H | + | - | ns | + | - | - | - | + | + | + | - | ns |
| G185V-H_VIN-H | ns* | - | + | + | - | ns | - | ns | + | + | - | ns |

* ns, not significant

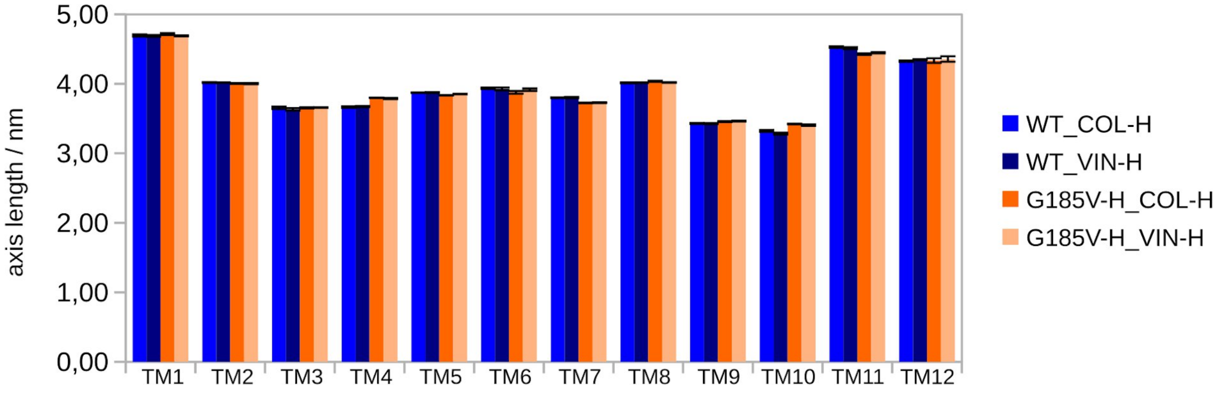


**Figure S3:** Comparison of individual transmembrane axis length for the WT and G185V variant (*holo* systems).

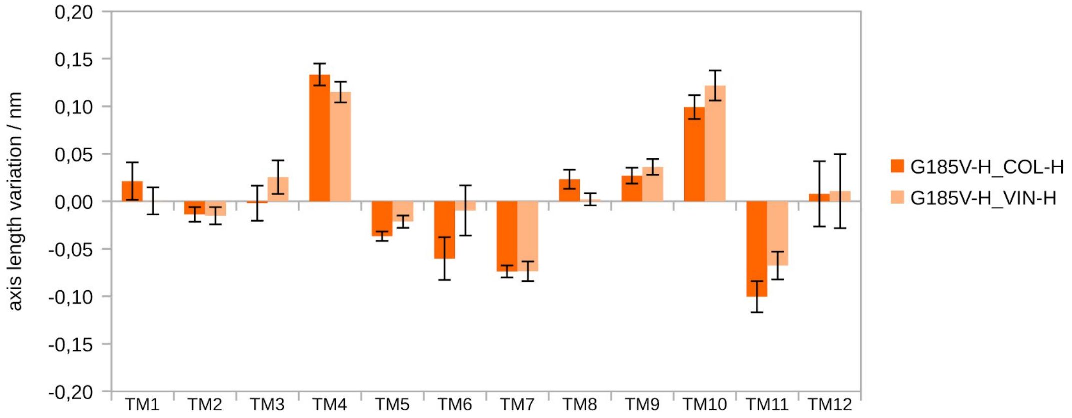


**Figure S4:** Variation in the transmembrane axis length for the *holo* G185V variant (difference from the *holo* WT).

¶
¶
¶

**Table S3.** Variation in the *z*-shift of the axis mid-points observed in the *holo* G185V variant transmembrane helices (compared to *holo* WT; +, positive mean changes; –, negative mean changes).

| | TM1 | TM2 | TM3 | TM4 | TM5 | TM6 | TM7 | TM8 | TM9 | TM10 | TM11 | TM12 |
|---|---|---|---|---|---|---|---|---|---|---|---|---|
| G185V-H_COL-H | – | + | ns* | – | ns | – | + | – | + | + | + | – |
| G185V-H_VIN-H | – | + | ns | – | ns | – | + | – | + | + | + | – |

* ns, not significant

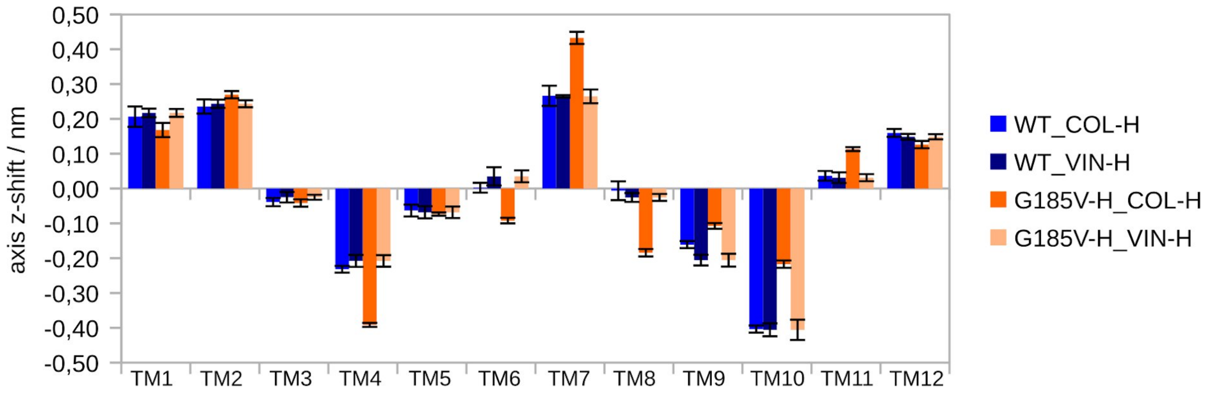


**Figure S5:** Comparison of individual transmembrane z-shift of the axis mid-points for the WT and G185V variant (*holo* systems).

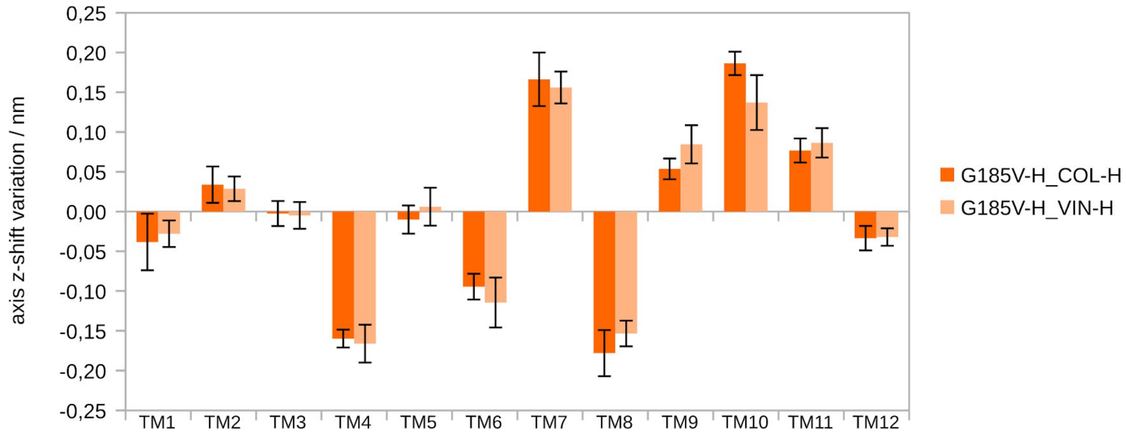


**Figure S6:** Variation in the transmembrane *z*-shift of the axis mid-points for the *holo* G185V variant (difference from the *holo* WT).

**Table S4.** Variation in the the axis total tilt observed in the *holo* G185V variant transmembrane helices (compared to the *holo* WT; +, positive mean changes; –, negative mean changes).

| | TM1 | TM2 | TM3 | TM4 | TM5 | TM6 | TM7 | TM8 | TM9 | TM10 | TM11 | TM12 |
|---|---|---|---|---|---|---|---|---|---|---|---|---|
| G185V-H_COL-H | ns* | – | + | + | + | + | ns | + | + | + | ns | – |
| G185V-H_VIN-H | ns | – | + | + | + | + | ns | + | + | + | ns | – |

* ns, not significant

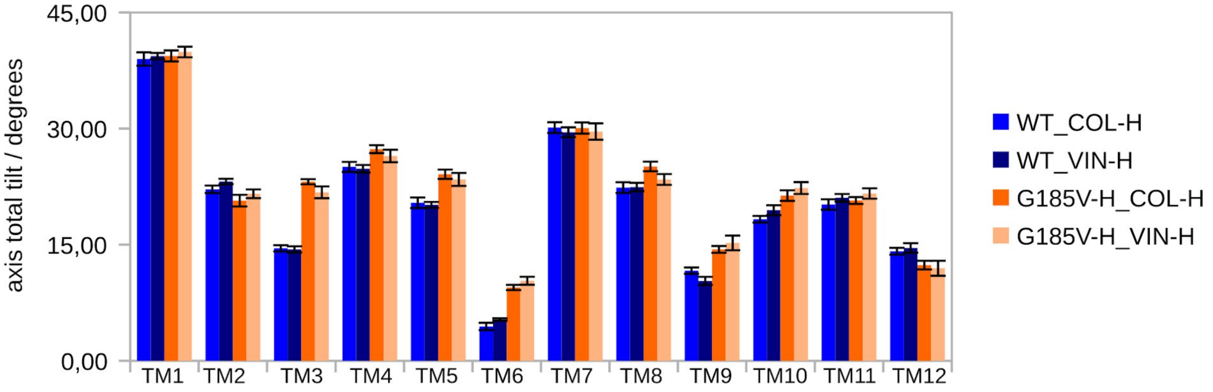


**Figure S7:** Comparison of individual transmembrane axis total tilt for the WT and G185V variant (*holo* systems).

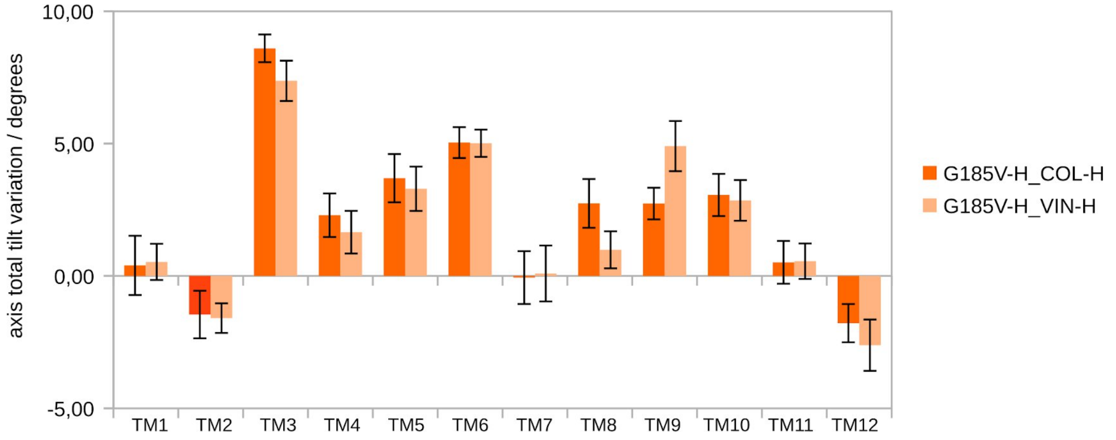


**Figure S8:** Variation in the transmembrane axis total tilt for the *holo* G185V variant (difference from the *holo* WT).

**Table S5.** Variation in the axis lateral tilt observed in the *holo* G185V variant transmembrane helices (compared to the *holo* WT; +, positive mean changes; –, negative mean changes).

| | TM1 | TM2 | TM3 | TM4 | TM5 | TM6 | TM7 | TM8 | TM9 | TM10 | TM11 | TM12 |
|---|---|---|---|---|---|---|---|---|---|---|---|---|
| G185V-H_COL-H | + | + | + | – | – | + | – | – | + | – | + | ns |
| G185V-H_VIN-H | + | + | + | ns* | + | + | – | ns | + | – | + | ns |

* ns, not significant

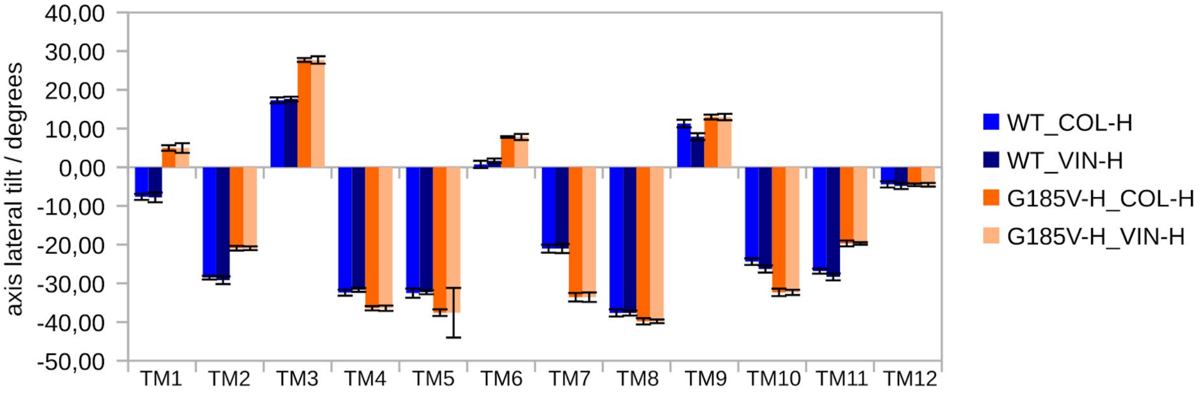


**Figure S9:** Comparison of individual transmembrane axis lateral tilt for the WT and G185V variant (*holo* systems).

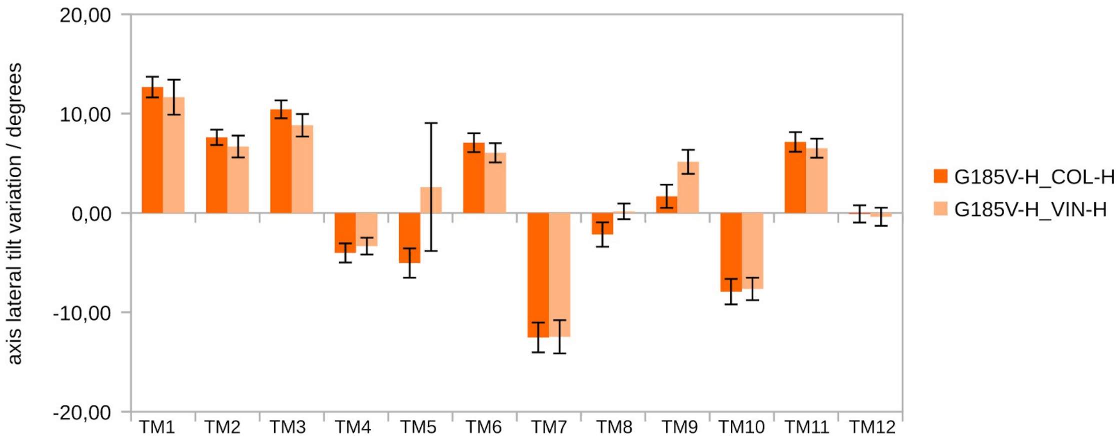


**Figure S10:** Variation in the transmembrane axis lateral tilt for the *holo* G185V variant (difference from the *holo* WT).

**Table S6.** Variation in the axis radial tilt observed in the *holo* G185V variant transmembrane helices (compared to the *holo* WT; +, positive mean changes; –, negative changes).

| | TM1 | TM2 | TM3 | TM4 | TM5 | TM6 | TM7 | TM8 | TM9 | TM10 | TM11 | TM12 |
|---|---|---|---|---|---|---|---|---|---|---|---|---|
| G185V-H_COL-H | ns* | – | + | – | + | – | + | – | – | – | – | + |
| G185V-H_VIN-H | ns | – | + | – | + | – | + | – | – | – | – | + |

* ns, not significant

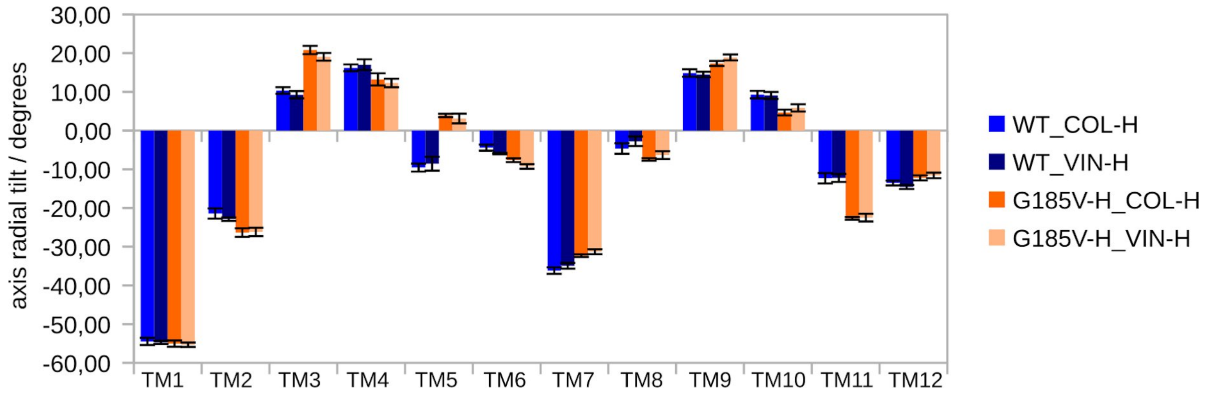


**Figure S11:** Comparison of individual transmembrane axis radial tilt for the WT and G185V variant (*holo* systems).

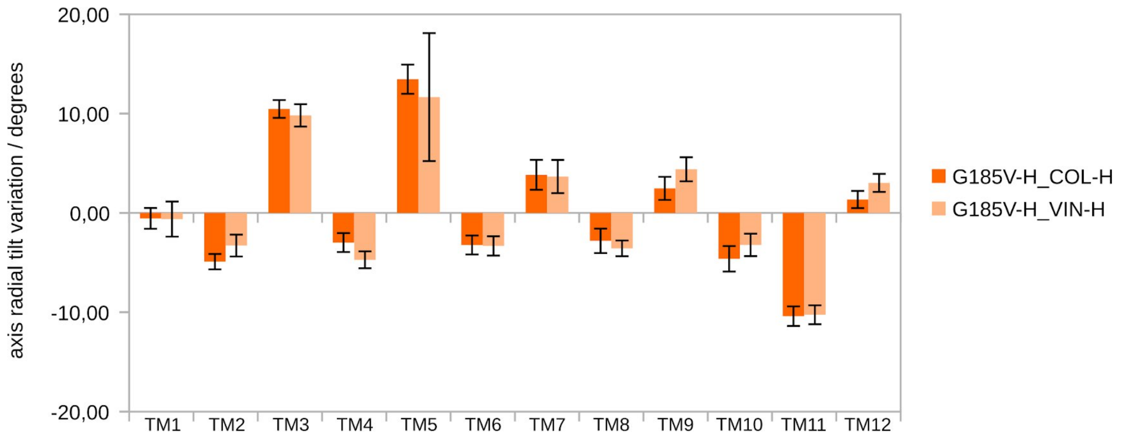


**Figure S12:** Variation in the transmembrane axis radial tilt for the *holo* G185V variant (difference from the *holo* WT).

**Table S7.** Variation in the axis distance observed in the *holo* G830V variant transmembrane helices (compared to the *holo* WT; +, positive mean changes; –, negative mean changes).

| | TM1 | TM2 | TM3 | TM4 | TM5 | TM6 | TM7 | TM8 | TM9 | TM10 | TM11 | TM12 |
|---|---|---|---|---|---|---|---|---|---|---|---|---|
| G830V-R_ACT-R | ns* | – | ns | ns | ns | + | ns | – | + | + | – | + |
| G830V-R_DOX-R | ns | – | ns | ns | – | + | + | – | ns | + | – | – |

* ns, not significant

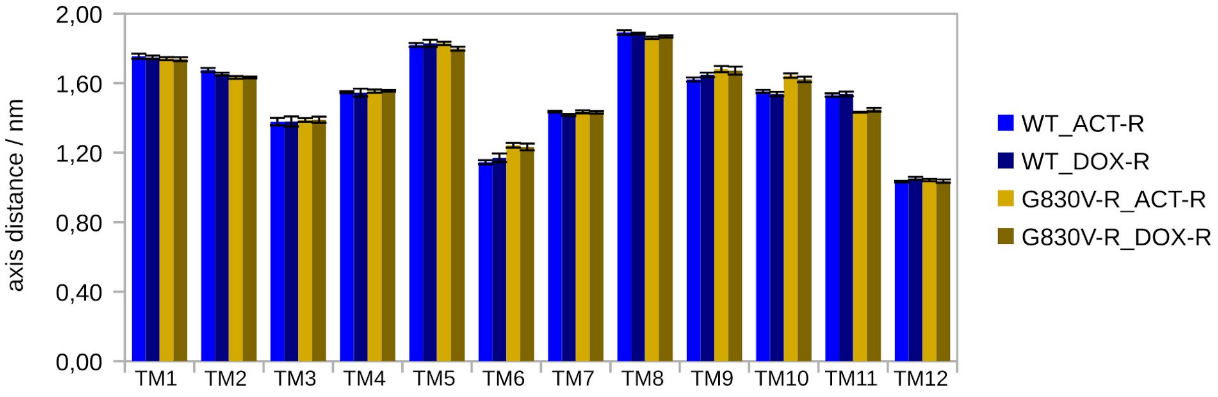


**Figure S13:** Comparison of individual transmembrane axis distance for the WT and G830V variant (*holo* systems).

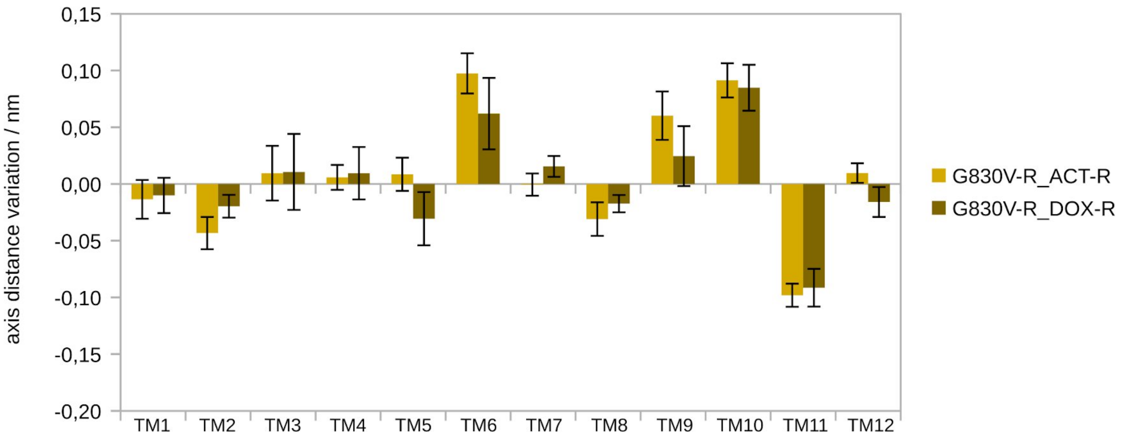


**Figure S14:** Variation in the transmembrane axis distance for the *holo* G830V variant (difference from the *holo* WT).

**Table S8.** Variation in the axis length observed in the *holo* G830V variant transmembrane helices (compared to the *holo* WT; +, positive mean changes; –, negative mean changes).

| | TM1 | TM2 | TM3 | TM4 | TM5 | TM6 | TM7 | TM8 | TM9 | TM10 | TM11 | TM12 |
|---|---|---|---|---|---|---|---|---|---|---|---|---|
| G830V-R_ACT-R | + | – | – | + | ns | – | + | + | + | + | – | + |
| G830V-R_DOX-R | ns* | ns | – | ns | – | – | ns | ns | + | + | – | + |

* ns, not significant

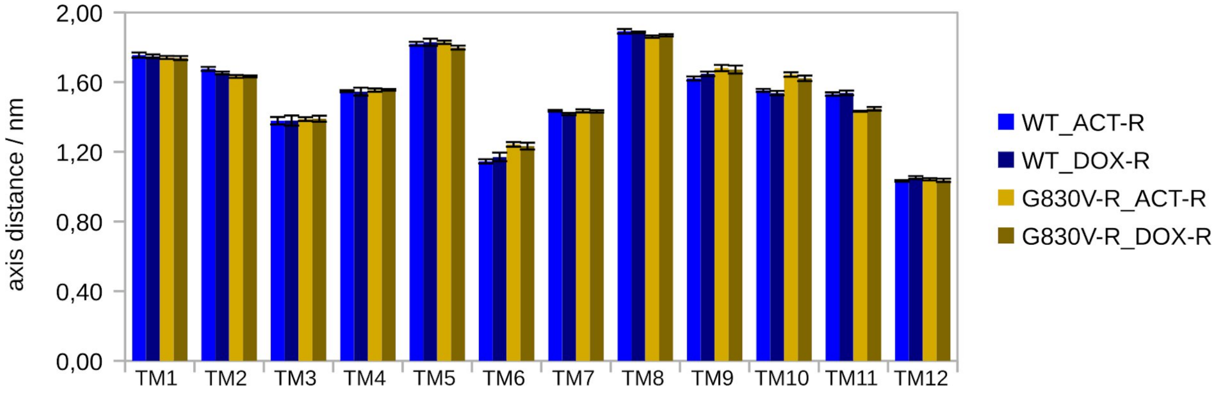


**Figure S15:** Comparison of individual transmembrane axis length for the WT and G830V variant (*holo* systems).

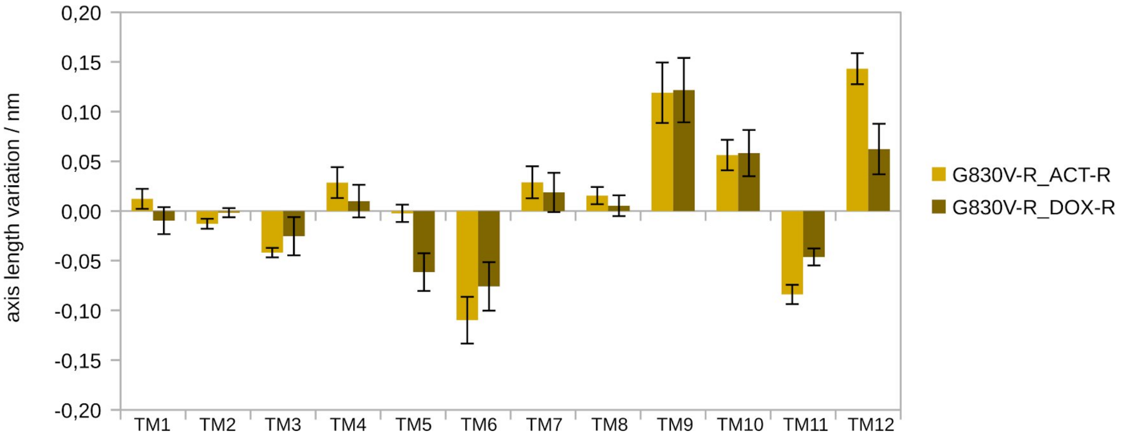


**Figure S16:** Variation in the transmembrane axis length for the *holo* G830V variant (difference from the *holo* WT).

**Table S9.** Variation in the *z*-shift of the axis mid-points observed in the *holo* G830V variant transmembrane helices (compared to the *holo* WT; +, positive mean changes; –, negative mean changes).

| | TM1 | TM2 | TM3 | TM4 | TM5 | TM6 | TM7 | TM8 | TM9 | TM10 | TM11 | TM12 |
|---|---|---|---|---|---|---|---|---|---|---|---|---|
| G830V-R_ACT-R | – | – | – | – | + | ns | + | + | + | + | ns | ns |
| G830V-R_DOX-R | – | ns* | – | – | – | – | + | + | + | + | + | + |

* ns, not significant

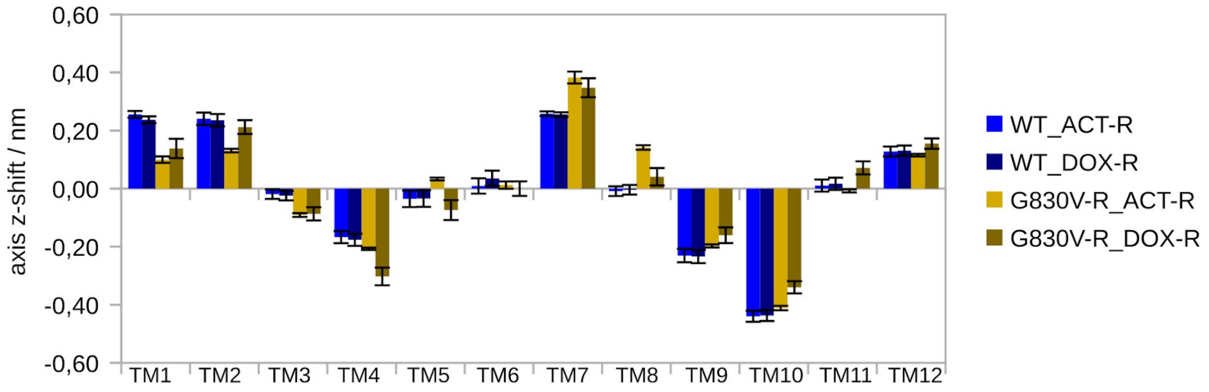


**Figure S17:** Comparison of individual transmembrane *z*-shift of the axis mid-points for the WT and G830V variant (*holo* systems).

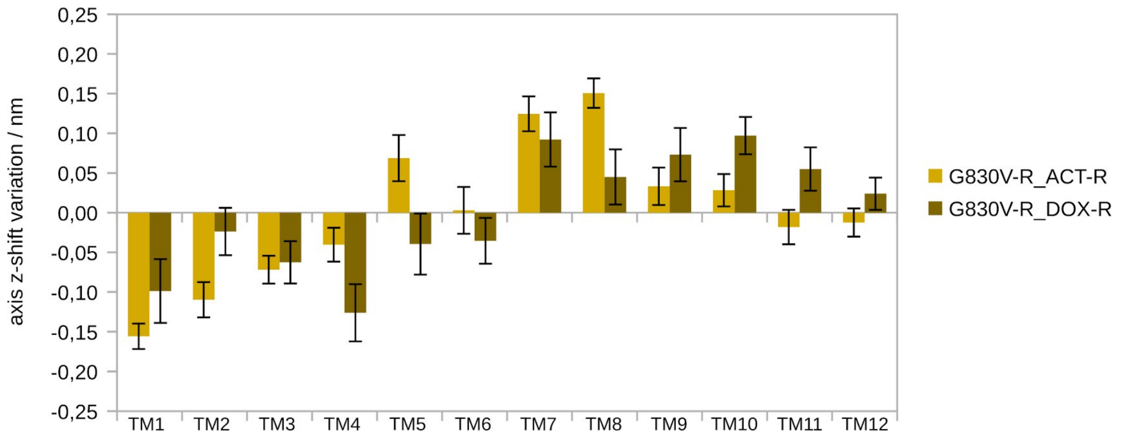


**Figure S18:** Variation in the transmembrane *z*-shift of the axis mid-points for the *holo* G830V variant (difference from the *holo* WT).

**Table S10.** Variation in the the axis total tilt observed in the *holo* G830V variant transmembrane helices (compared to the *holo* WT; +, positive mean changes; –, negative mean changes).

| | TM1 | TM2 | TM3 | TM4 | TM5 | TM6 | TM7 | TM8 | TM9 | TM10 | TM11 | TM12 |
|---|---|---|---|---|---|---|---|---|---|---|---|---|
| G830V-R_ACT-R | – | – | + | + | + | – | + | + | + | – | – | – |
| G830V-R_DOX-R | – | – | + | + | + | – | + | + | + | ns* | – | – |

* ns, not significant

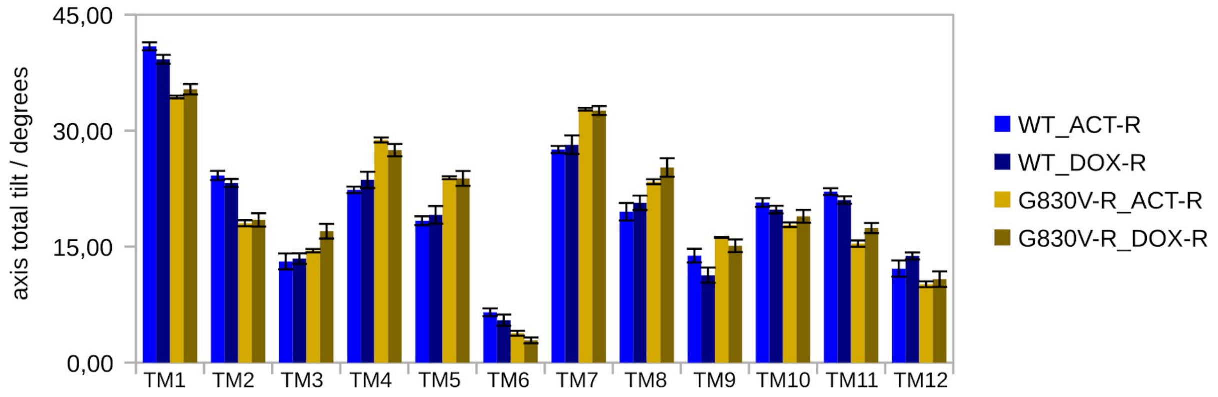


**Figure S19:** Comparison of individual transmembrane axis total tilt for the WT and *holo* G830V variant (*holo* systems).

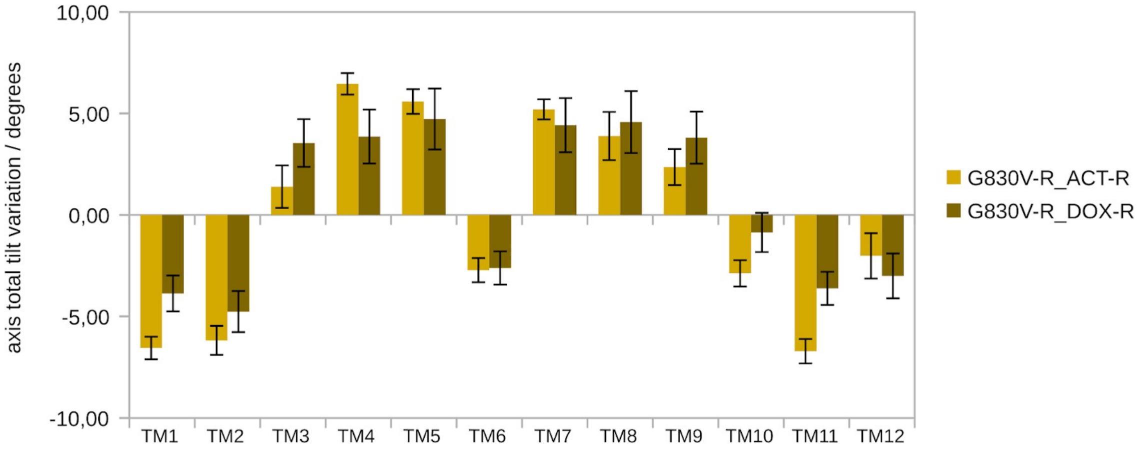


**Figure S20:** Variation in the transmembrane axis total tilt for the *holo* G830V variant (difference from the *holo* WT).

**Table S11.** Variation in the axis lateral tilt observed in the *holo* G830V variant transmembrane helices (compared to the *holo* WT; +, positive mean changes; –, negative mean changes).

| | TM1 | TM2 | TM3 | TM4 | TM5 | TM6 | TM7 | TM8 | TM9 | TM10 | TM11 | TM12 |
|---|---|---|---|---|---|---|---|---|---|---|---|---|
| G830V-R_ACT-R | + | + | – | – | – | – | – | – | + | + | + | + |
| G830V-R_DOX-R | + | + | + | – | – | ns* | – | – | + | – | + | + |

* ns, not significant

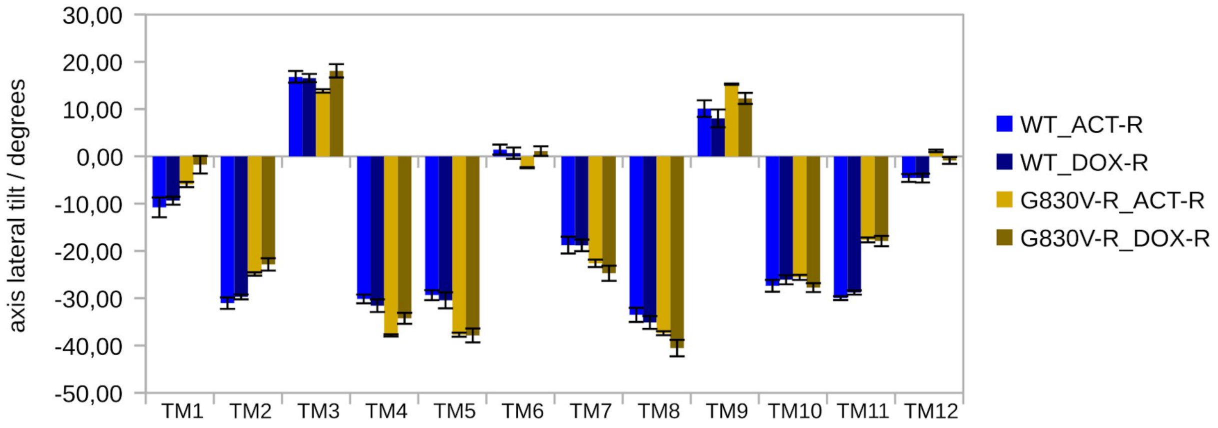


**Figure S21:** Comparison of individual transmembrane axis lateral tilt for the WT and G830V variant (*holo* systems).

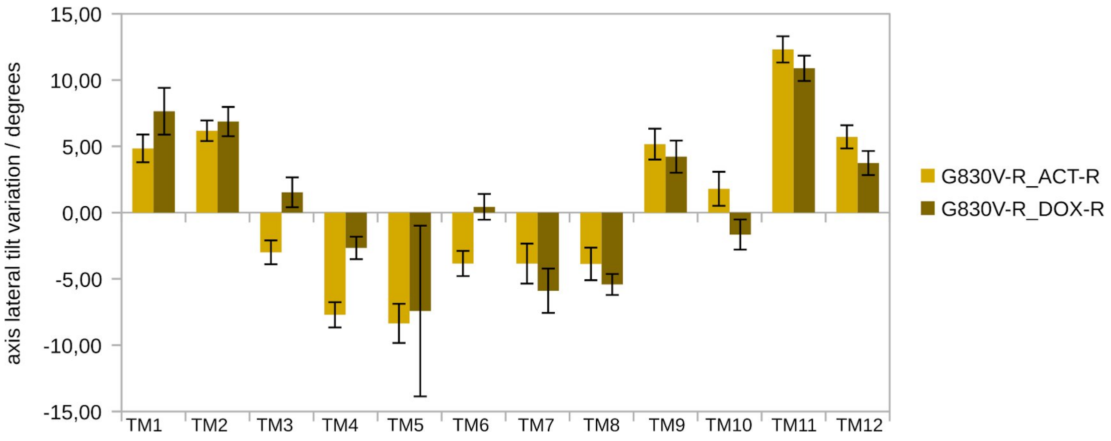


¶

**Figure S22:** Variation in the transmembrane axis lateral tilt for the *holo* G830V variant (difference from the *holo* WT).

**Table S12.** Variation in the axis radial tilt observed in the *holo* G830V variant transmembrane helices (compared to the *holo* WT; +, positive mean changes; –, negative mean changes).

| | TM1 | TM2 | TM3 | TM4 | TM5 | TM6 | TM7 | TM8 | TM9 | TM10 | TM11 | TM12 |
|---|---|---|---|---|---|---|---|---|---|---|---|---|
| G830V-R_ACT-R | + | + | + | + | ns* | + | – | – | + | ns | ns | + |
| G830V-R_DOX-R | + | + | + | + | + | + | – | – | + | – | – | + |

* ns, not significant

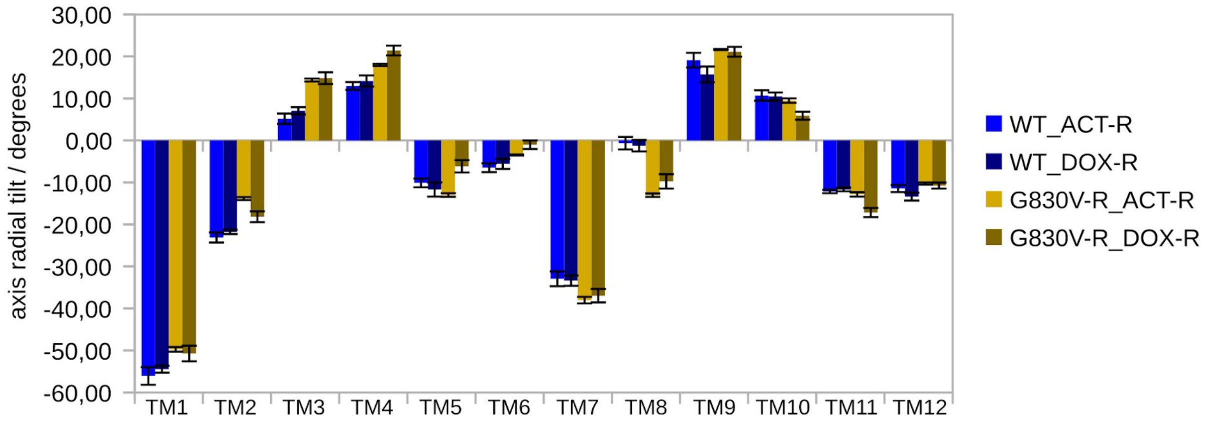


**Figure S23:** Comparison of individual transmembrane axis radial tilt for the WT and G830V varian (*holo* systems).

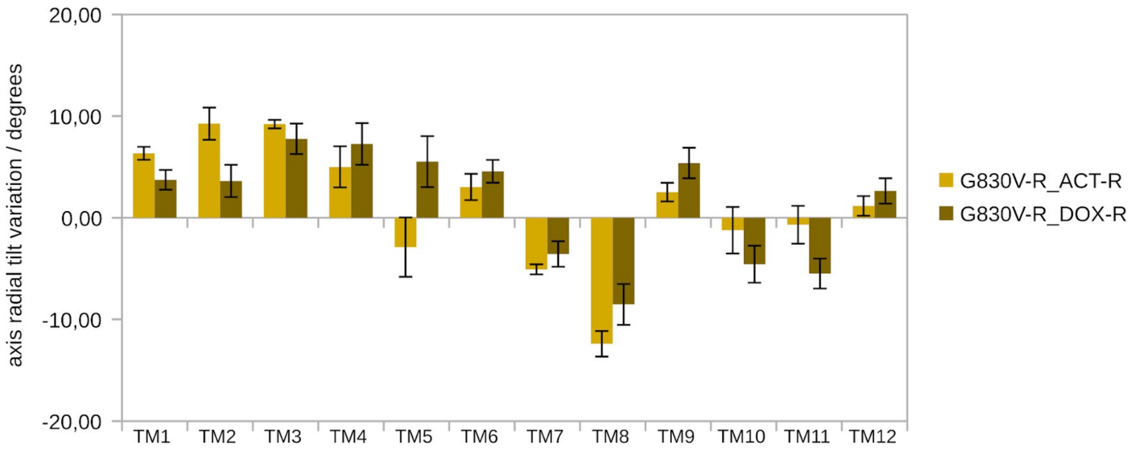


**Figure S24:** Variation in the transmembrane axis radial tilt for the *holo* G830V variant (difference from the *holo* WT).

**Table S13.** Variation in the axis distance observed in the *holo* F978A variant transmembrane helices (compared to the *holo* WT; +, positive mean changes; –, negative mean changes).

| | TM1 | TM2 | TM3 | TM4 | TM5 | TM6 | TM7 | TM8 | TM9 | TM10 | TM11 | TM12 |
|---|---|---|---|---|---|---|---|---|---|---|---|---|
| F978A-M_ACT-R | ns* | – | – | – | – | + | + | ns | – | + | + | + |
| F978A-M_CYC-M | + | – | ns | + | + | + | ns | – | – | + | + | + |
| F978A-M_VIN-H | ns | ns | – | – | ns | ns | + | ns | – | + | + | + |

* ns, not significant

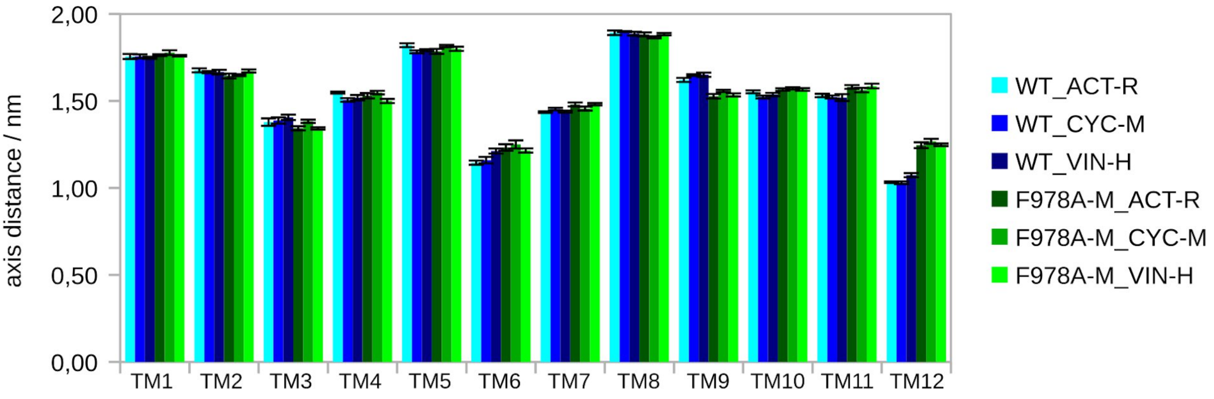


**Figure S25:** Comparison of individual transmembrane axis distance for the WT and F978A variant (*holo* systems).

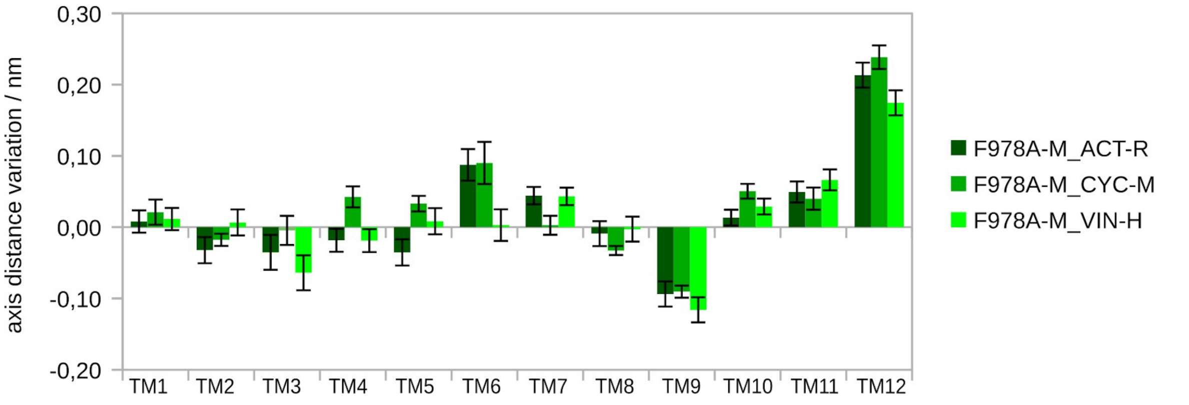


**Figure S26:** Variation in the transmembrane axis distance for the *holo* F978A variant (difference from the *holo* WT).

**Table S14.** Variation in the axis length observed in the *holo* F978A variant transmembrane helices (compared to the *holo* WT; +, positive mean changes; –, negative mean changes).

| | TM1 | TM2 | TM3 | TM4 | TM5 | TM6 | TM7 | TM8 | TM9 | TM10 | TM11 | TM12 |
|---|---|---|---|---|---|---|---|---|---|---|---|---|
| F978A-M_ACT-R | + | + | – | + | – | – | + | + | – | + | – | – |
| F978A-M_CYC-M | – | ns | – | + | ns | – | + | + | – | + | – | – |
| F978A-M_VIN-H | ns* | + | – | ns | – | – | + | ns | – | + | – | – |

* ns, not significant

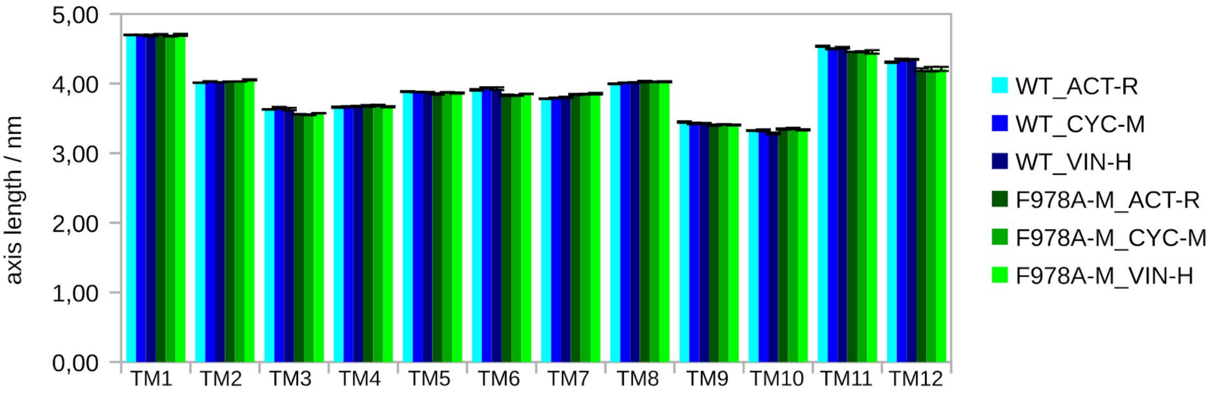


**Figure S27:** Comparison of individual transmembrane axis length for the WT and F978A variant (*holo* systems).

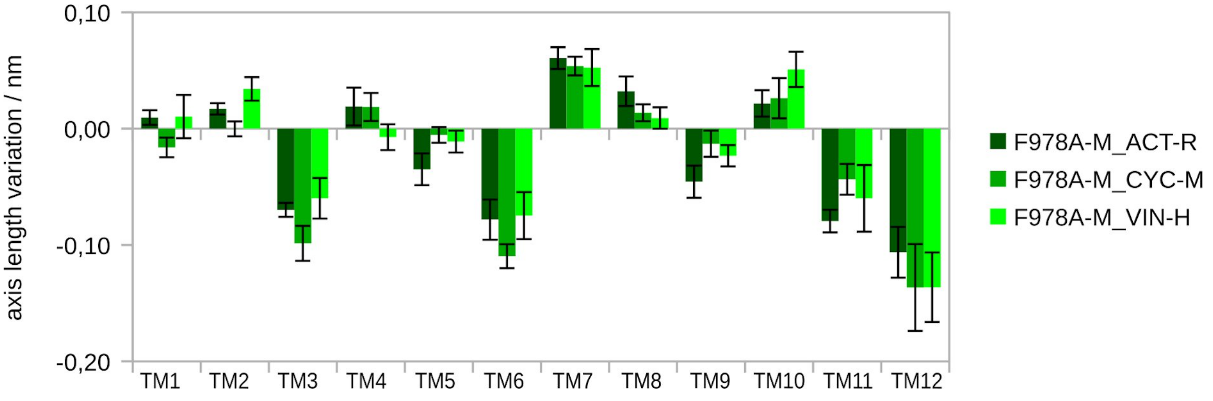


**Figure S28:** Variation in the transmembrane axis length for the *holo* F978A variant (difference from the *holo* WT).

**Table S15.** Variation in the *z*-shift of the axis mid-points observed in the *holo* F978A variant transmembrane helices (compared to the *holo* WT; +, positive mean changes; –, negative changes).

| | TM1 | TM2 | TM3 | TM4 | TM5 | TM6 | TM7 | TM8 | TM9 | TM10 | TM11 | TM12 |
|---|---|---|---|---|---|---|---|---|---|---|---|---|
| F978A-M_ACT-R | – | – | – | – | – | – | + | ns | + | + | ns | + |
| F978A-M_CYC-M | – | ns | – | – | – | – | + | + | + | + | ns | + |
| F978A-M_VIN-H | ns* | ns | – | – | – | – | + | ns | ns | + | ns | + |

* ns, not significant

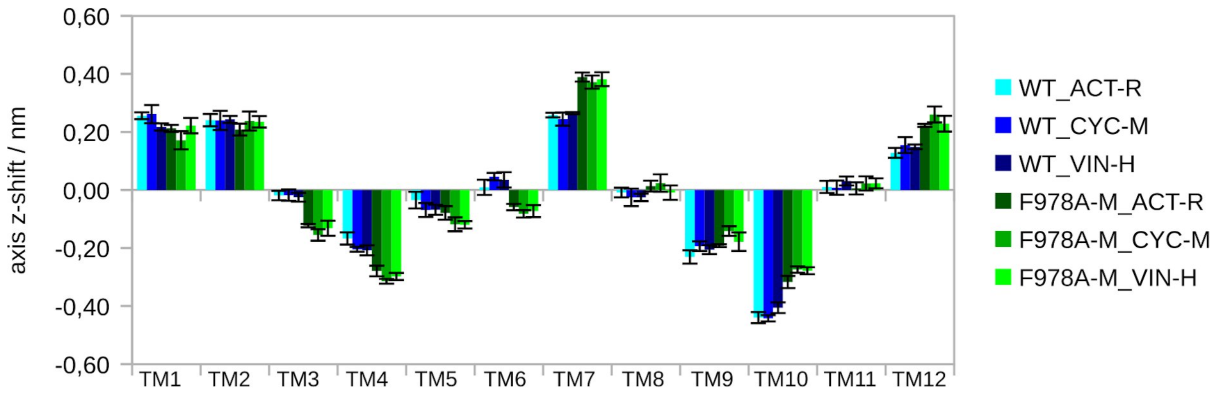


**Figure S29:** Comparison of individual transmembrane *z*-shift of the axis mid-points for the WT and F978A variant (*holo* systems).

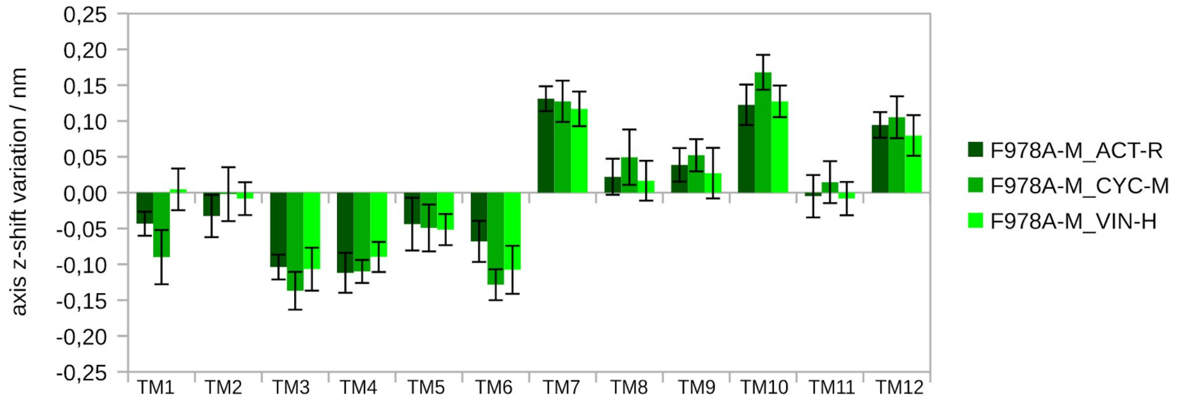


**Figure S30:** Variation in the transmembrane *z*-shift of the axis mid-points for the *holo* F978A variant (difference from the *holo* WT).

**Table S16.** Variation in the the axis total tilt observed in the *holo* F978A variant transmembrane helices (compared to the *holo* WT; +, positive mean changes; –, negative changes).

| | TM1 | TM2 | TM3 | TM4 | TM5 | TM6 | TM7 | TM8 | TM9 | TM10 | TM11 | TM12 |
|---|---|---|---|---|---|---|---|---|---|---|---|---|
| F978A-M_ACT-R | ns* | – | + | + | + | ns | + | + | – | – | – | + |
| F978A-M_CYC-M | – | – | + | + | + | ns | + | + | – | – | – | + |
| F978A-M_VIN-H | + | – | + | + | + | ns | ns | + | – | ns | – | + |

* ns, not significant

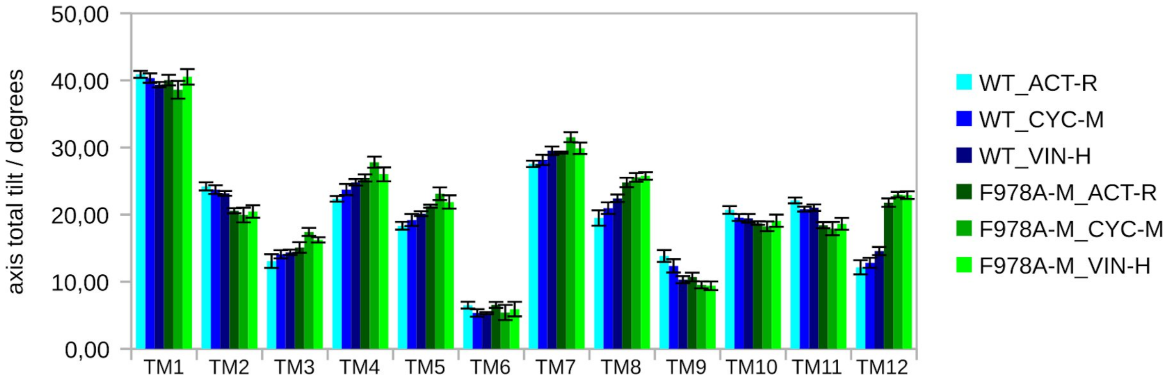


**Figure S31:** Comparison of individual transmembrane axis total tilt for the WT and F978A variant (*holo* systems).

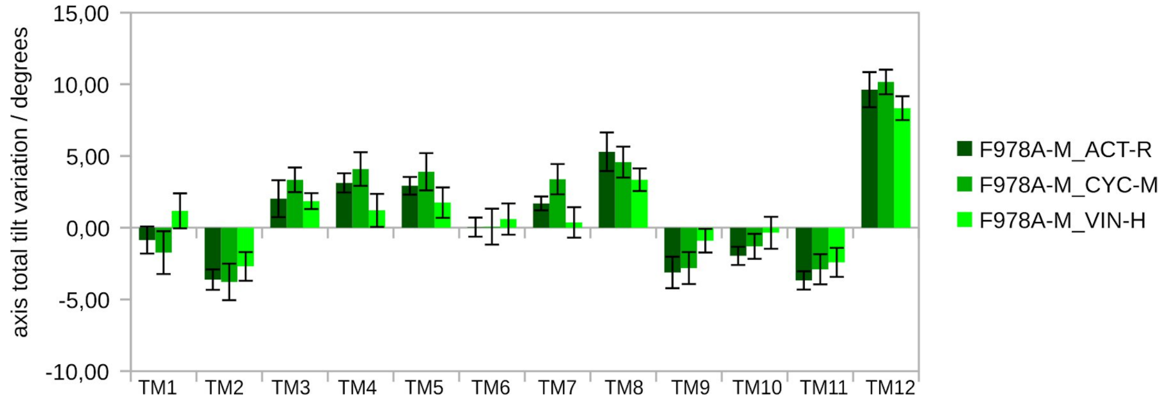


**Figure S32:** Variation in the transmembrane axis total tilt for the *holo* F978A variant (difference from the *holo* WT).

**Table S17.** Variation in the taxis lateral tilt observed in the *holo* F978A variant transmembrane helices (compared to the *holo* WT; +, positive mean changes; –, negative mean changes).

| | TM1 | TM2 | TM3 | TM4 | TM5 | TM6 | TM7 | TM8 | TM9 | TM10 | TM11 | TM12 |
|---|---|---|---|---|---|---|---|---|---|---|---|---|
| F978A-M_ACT-R | + | + | ns* | – | – | + | – | – | ns | + | + | ns |
| F978A-M_CYC-M | + | + | ns | ns | – | + | – | – | ns | + | + | ns |
| F978A-M_VIN-H | + | + | + | ns | – | + | – | – | ns | ns | + | ns |

* ns, not significant

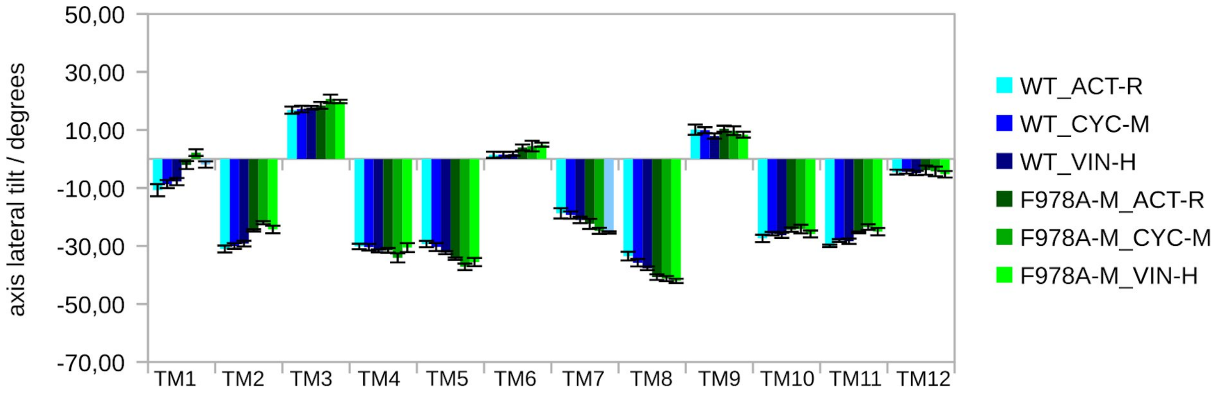


**Figure S33:** Comparison of individual transmembrane axis lateral tilt for the WT and F978A variant (*holo* systems).

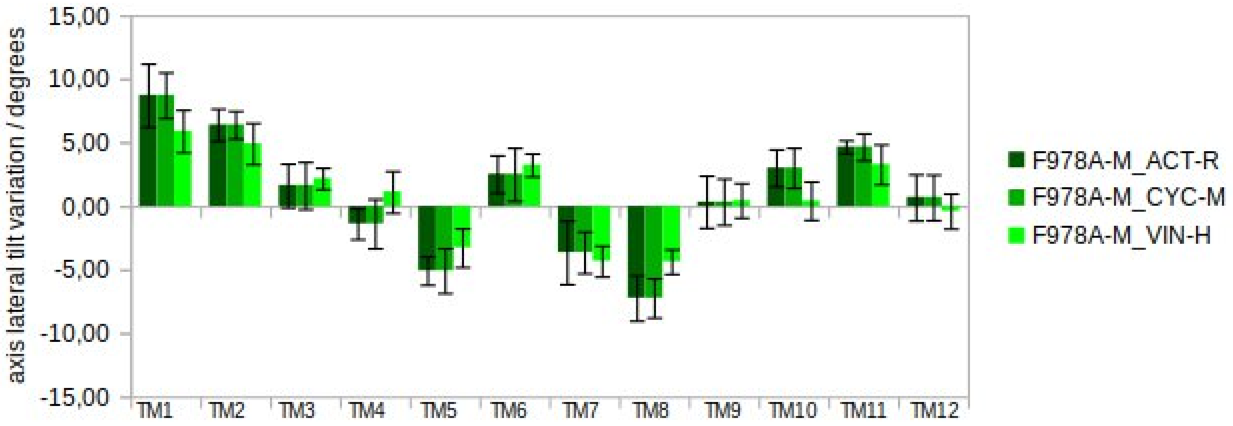


**Figure S34:** Variation in the transmembrane axis lateral tilt for the *holo* F978A variant (difference from the *holo* WT).

**Table S18.** Variation in the axis radial tilt observed in the *holo* F978A variant transmembrane helices (compared to the *holo* WT; +, positive mean changes; –, negative mean changes).

| | TM1 | TM2 | TM3 | TM4 | TM5 | TM6 | TM7 | TM8 | TM9 | TM10 | TM11 | TM12 |
|---|---|---|---|---|---|---|---|---|---|---|---|---|
| F978A-M_ACT-R | ns* | ns | + | + | + | ns | – | – | – | + | ns | – |
| F978A-M_CYC-M | ns | ns | + | + | + | + | – | – | – | + | – | – |
| F978A-M_VIN-H | ns | + | + | + | + | ns | ns | – | – | + | ns | – |

* ns, not significant

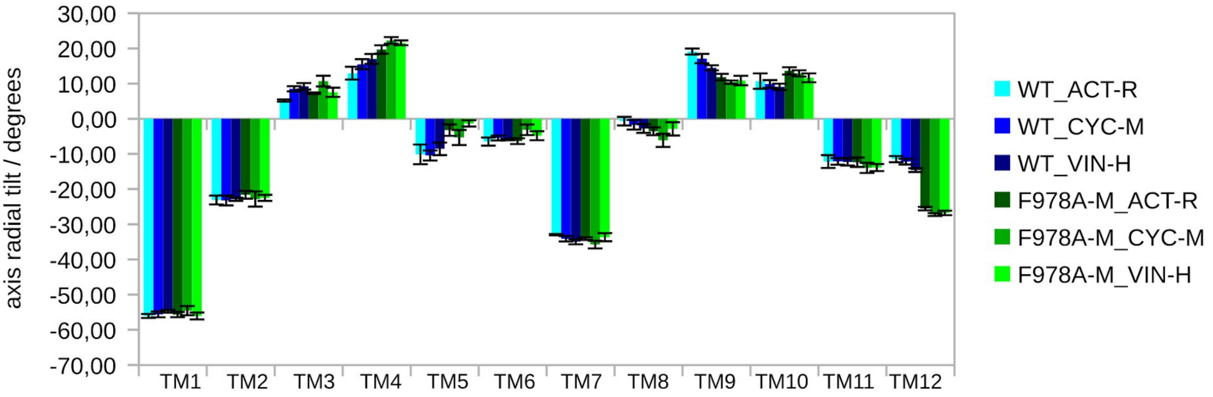


**Figure S35:** Comparison of individual transmembrane axis radial tilt for the WT and F978A variant (*holo* systems).

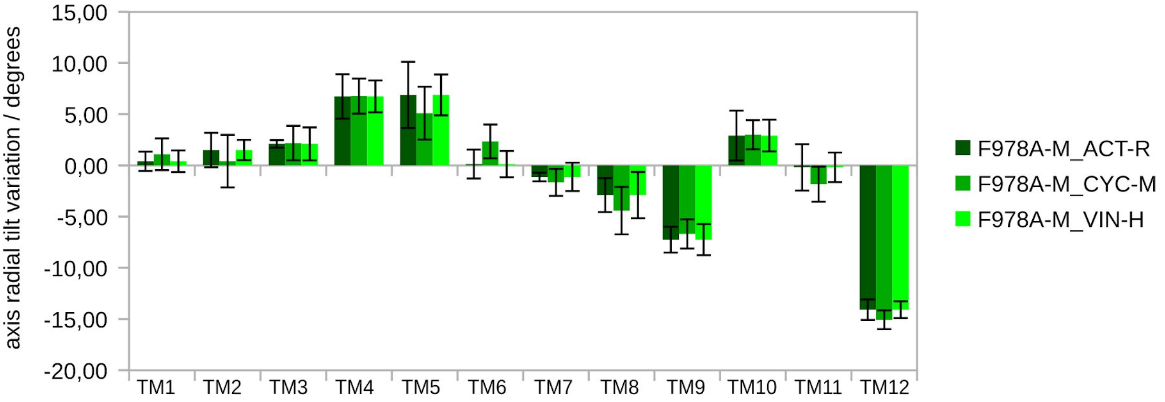


**Figure S36:** Variation in the transmembrane axis radial tilt for the *holo* F978A variant (difference from the *holo* WT).

**Table S19.** Variation in the axis distance observed in the *holo* ΔF335 variant transmembrane helices (compared to the *holo* WT; +, positive mean changes; –, negative mean changes).

| | TM1 | TM2 | TM3 | TM4 | TM5 | TM6 | TM7 | TM8 | TM9 | TM10 | TM11 | TM12 |
|---|---|---|---|---|---|---|---|---|---|---|---|---|
| ΔF335-M_CYC-M | + | + | + | ns | + | + | + | ns | – | ns | + | – |
| ΔF335-M_DOX-R | + | + | ns* | – | ns | + | + | ns | – | + | + | – |
| ΔF335-M_VLS-M | + | + | – | – | + | ns | ns | – | – | + | + | – |

* ns, not significant

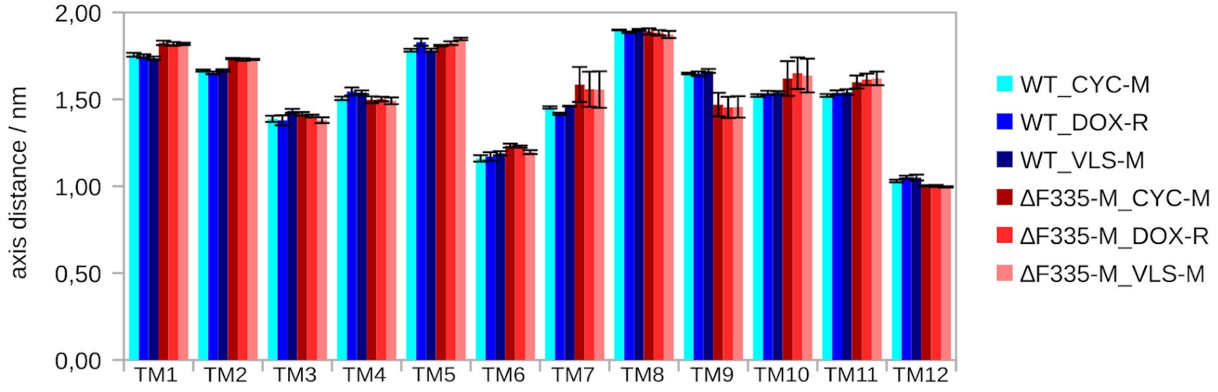


**Figure S37:** Comparison of individual transmembrane axis distance for the WT and ΔF335 variant (*holo* systems).

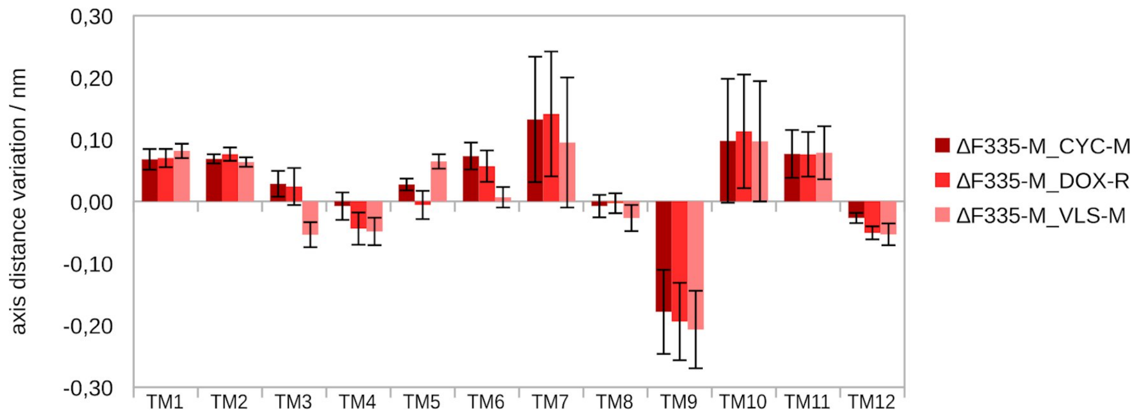


**Figure S38:** Variation in the transmembrane axis distance for the *holo* ΔF335 variant (difference from the *holo* WT).

**Table S20.** Variation in the axis length observed in the *holo* ΔF335 variant transmembrane helices (compared to the *holo* WT; +, positive mean changes; –, negative mean changes).

| | TM1 | TM2 | TM3 | TM4 | TM5 | TM6 | TM7 | TM8 | TM9 | TM10 | TM11 | TM12 |
|---|---|---|---|---|---|---|---|---|---|---|---|---|
| ΔF335-M_CYC-M | ns* | – | + | + | – | + | ns | ns | – | + | – | – |
| ΔF335-M_DOX-R | – | ns | + | + | – | + | ns | ns | – | + | – | – |
| ΔF335-M_VLS-M | – | – | + | + | – | + | – | ns | ns | + | – | – |

* ns, not significant

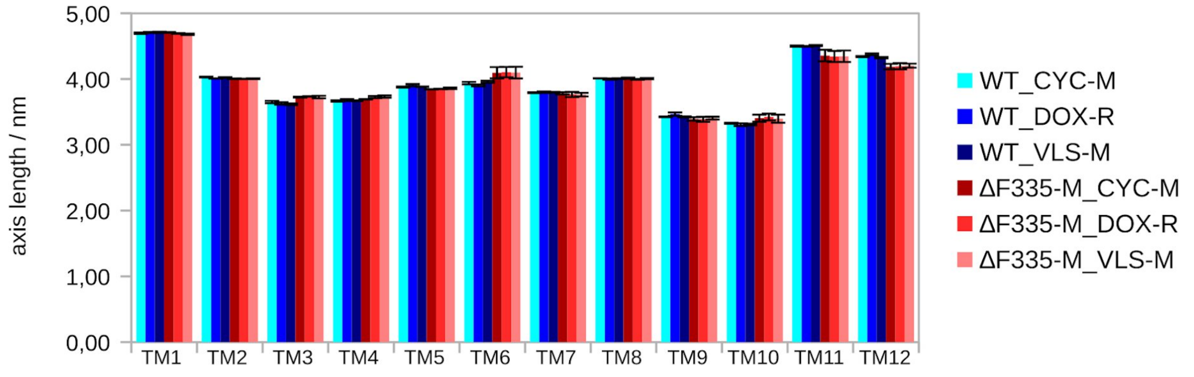


**Figure S39:** Comparison of individual transmembrane axis length for the WT and ΔF335 variant (*holo* systems).

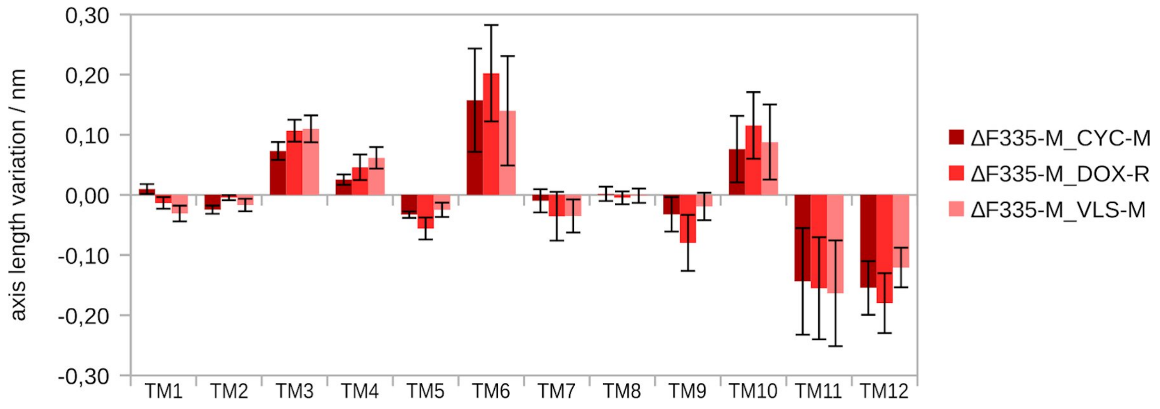


**Figure S40:** Variation in the transmembrane axis length for the *holo* ΔF335 variant (difference from the *holo* WT).

**Table S21.** Variation in the *z*-shift of the axis mid-points observed in the *holo* ΔF335 variant transmembrane helices (compared to the *holo* WT; +, positive mean changes; –, negative mean changes).

| | TM1 | TM2 | TM3 | TM4 | TM5 | TM6 | TM7 | TM8 | TM9 | TM10 | TM11 | TM12 |
|---|---|---|---|---|---|---|---|---|---|---|---|---|
| ΔF335-M_CYC-M | – | – | ns | ns | + | – | + | ns | + | ns | ns | ns |
| ΔF335-M_DOX-R | ns* | – | + | – | ns | ns | + | – | + | ns | ns | ns |
| ΔF335-M_VLS-M | – | – | + | ns | ns | ns | + | ns | + | – | ns | ns |

* ns, not significant

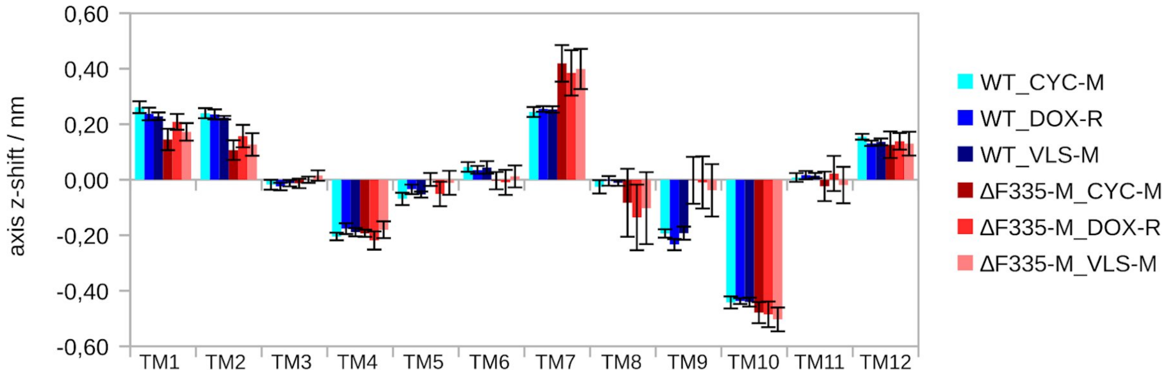


**Figure S41:** Comparison of individual transmembrane *z*-shift of the axis mid-points for the WT and ΔF335 variant (*holo* systems).

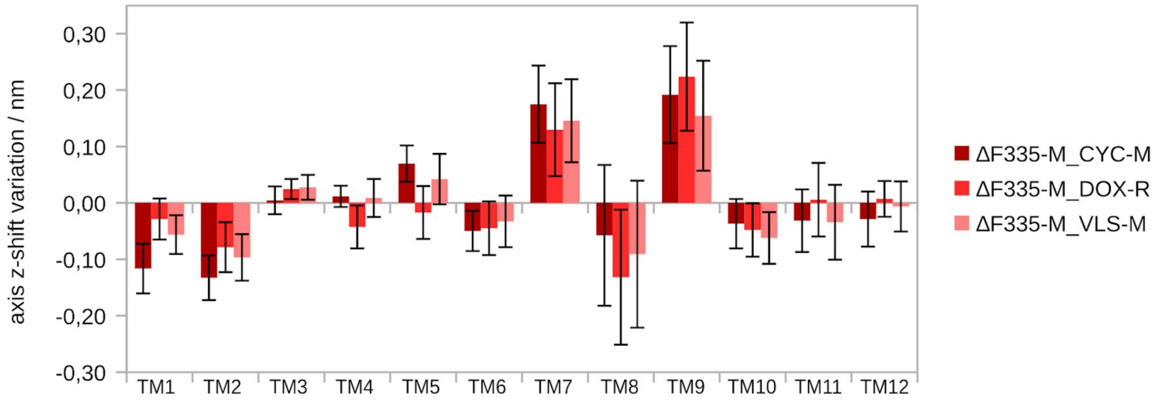


**Figure S42:** Variation in the transmembrane *z*-shift of the axis mid-points for the *holo* ΔF335 variant (difference from the *holo* WT).

**Table S22.** Variation in the the axis total tilt observed in the *holo* ΔF335 variant transmembrane helices (compared to the *holo* WT; +, positive mean changes; –, negative mean changes).

| | TM1 | TM2 | TM3 | TM4 | TM5 | TM6 | TM7 | TM8 | TM9 | TM10 | TM11 | TM12 |
|---|---|---|---|---|---|---|---|---|---|---|---|---|
| ΔF335-M_CYC-M | – | – | – | ns | + | ns | + | ns | – | ns | ns | + |
| ΔF335-M_DOX-R | – | – | ns* | ns | + | + | + | ns | – | ns | ns | ns |
| ΔF335-M_VLS-M | – | – | ns | ns | + | ns | + | – | – | ns | ns | ns |

* ns, not significant

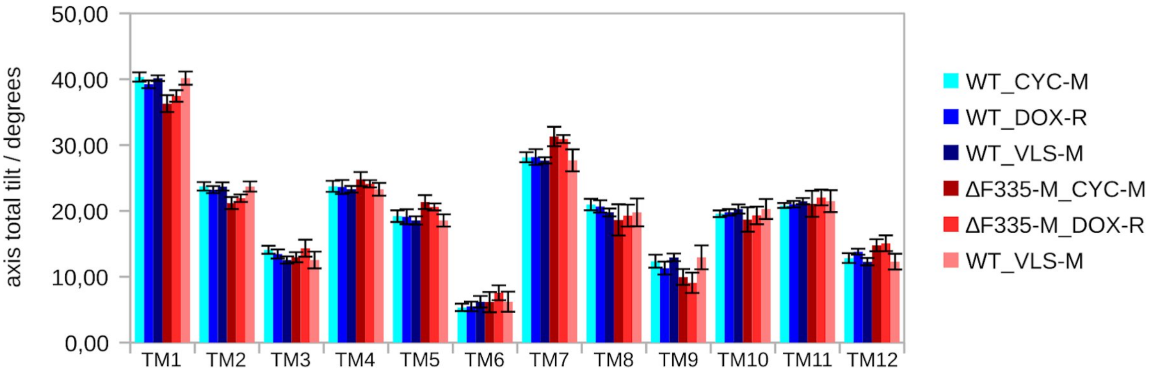


**Figure S43:** Comparison of individual transmembrane axis total tilt for the WT and ΔF335 variant (*holo* systems).

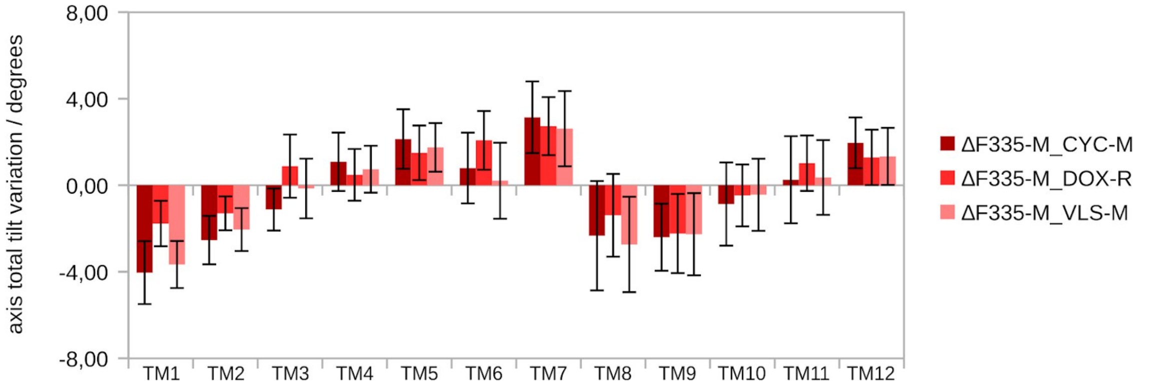


**Figure S44:** Variation in the transmembrane axis total tilt for the *holo* ΔF335 variant (difference from the *holo* WT).

**Table S23.** Variation in the taxis lateral tilt observed in the *holo* ΔF335 variant transmembrane helices (compared to the *holo* WT; +, positive mean changes; –, negative mean changes).

| | TM1 | TM2 | TM3 | TM4 | TM5 | TM6 | TM7 | TM8 | TM9 | TM10 | TM11 | TM12 |
|---|---|---|---|---|---|---|---|---|---|---|---|---|
| ΔF335-M_CYC-M | + | ns | ns | ns | – | + | – | + | ns | ns | ns | ns |
| ΔF335-M_DOX-R | + | ns | ns | + | – | + | – | + | ns | ns | ns | ns |
| ΔF335-M_VLS-M | ns* | ns | ns | ns | – | + | ns | + | ns | ns | ns | ns |

* ns, not significant

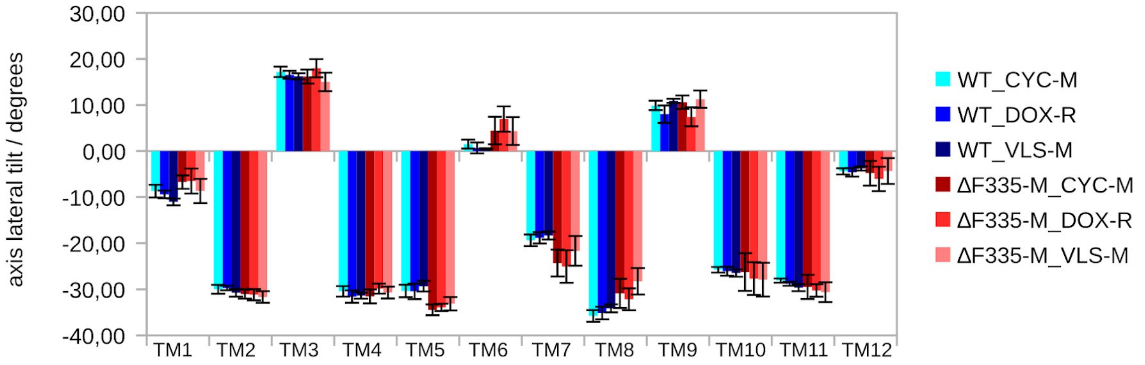


**Figure S45:** Comparison of individual transmembrane axis lateral tilt for the WT and ΔF335 variant (*holo* systems).

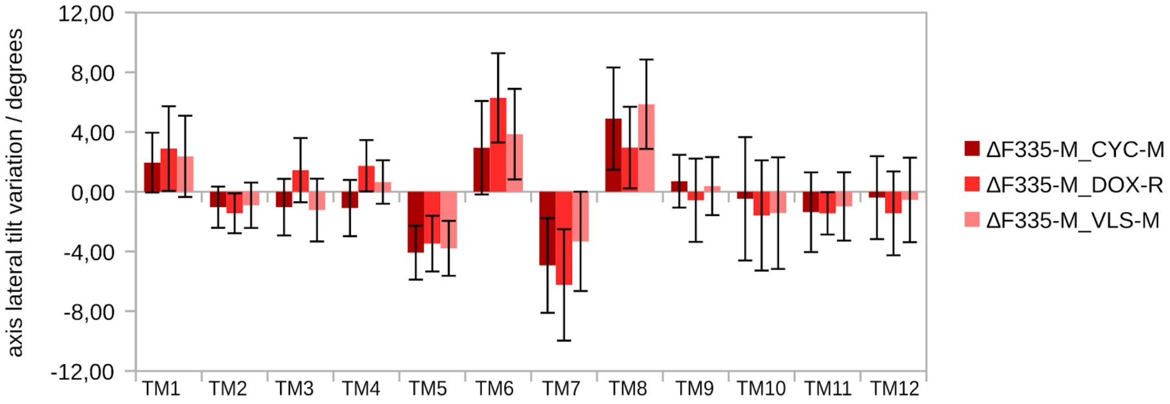


**Figure S46:** Variation in the transmembrane axis lateral tilt for the *holo* ΔF335 variant (difference from the *holo* WT).

**Table S24.** Variation in the axis radial tilt observed in the *holo* ΔF335 variant transmembrane helices (compared to the *holo* WT; +, positive mean changes; –, negative mean changes).

| | TM1 | TM2 | TM3 | TM4 | TM5 | TM6 | TM7 | TM8 | TM9 | TM10 | TM11 | TM12 |
|---|---|---|---|---|---|---|---|---|---|---|---|---|
| ΔF335-M_CYC-M | + | + | ns | ns | + | + | – | – | – | ns | ns | ns |
| ΔF335-M_DOX-R | ns* | + | ns | + | + | + | – | – | – | + | ns | ns |
| ΔF335-M_VLS-M | + | + | ns | ns | ns | + | – | – | – | + | ns | ns |

* ns, not significant

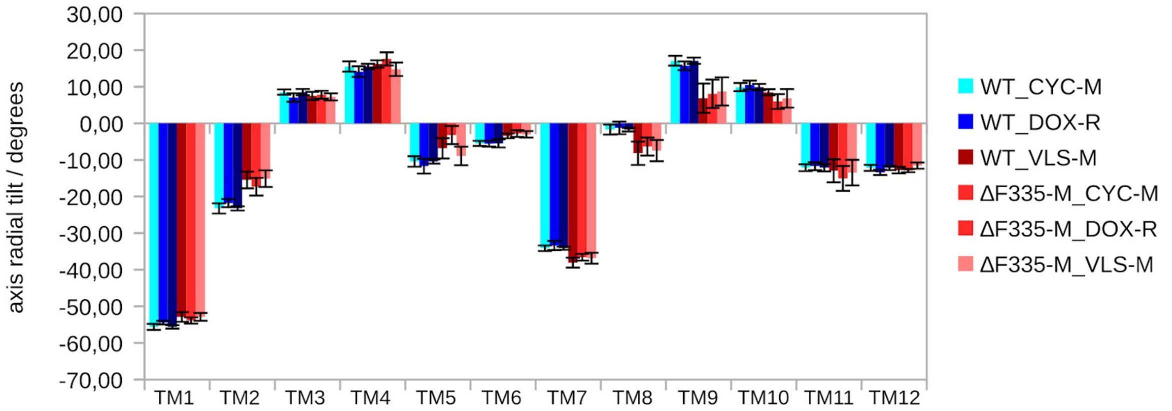


**Figure S47:** Comparison of individual transmembrane axis radial tilt for the WT and ΔF335 variant (*holo* systems).

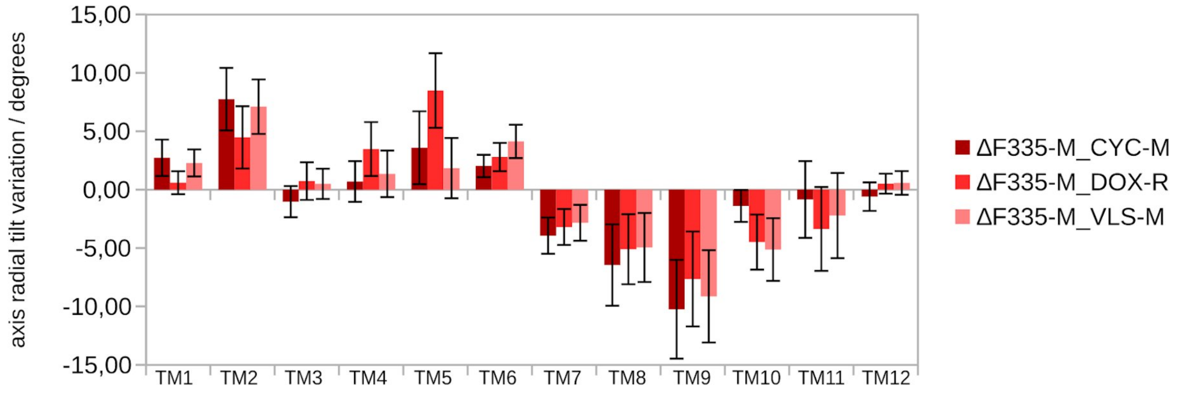


**Figure S48:** Variation in the transmembrane axis radial tilt for the WT and ΔF335 variant (difference from the *holo* WT).

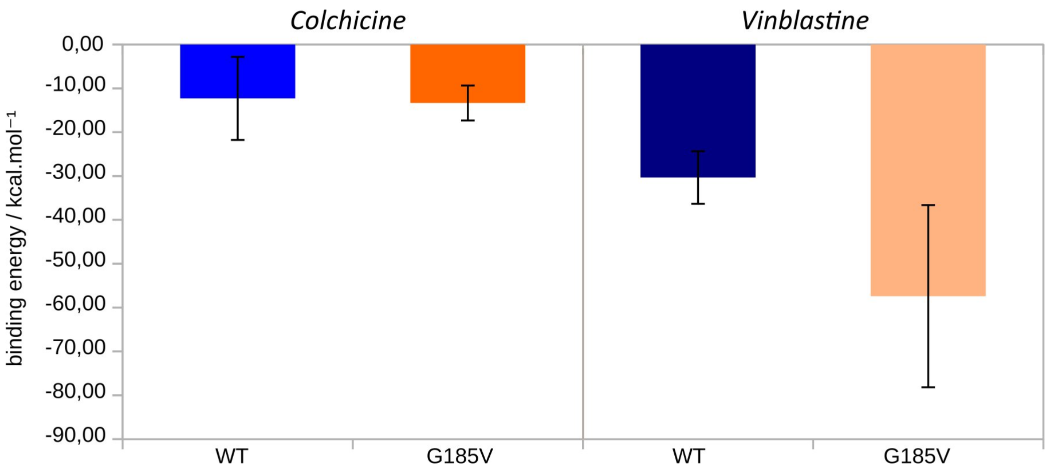


**Figure S49:** Relative Free-energies of binding ($\Delta G_{bind}$) for colchicine and vinblastine in the WT and G185V variant.

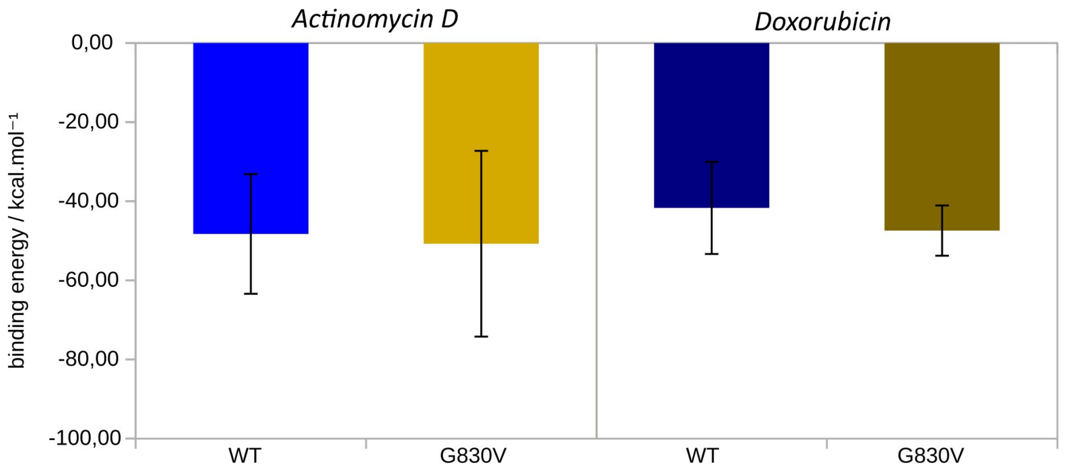


**Figure S50:** Relative Free energies of binding ($\Delta G_{bind}$) for actinomycin D and doxorubicin in the WT and G830V variant.

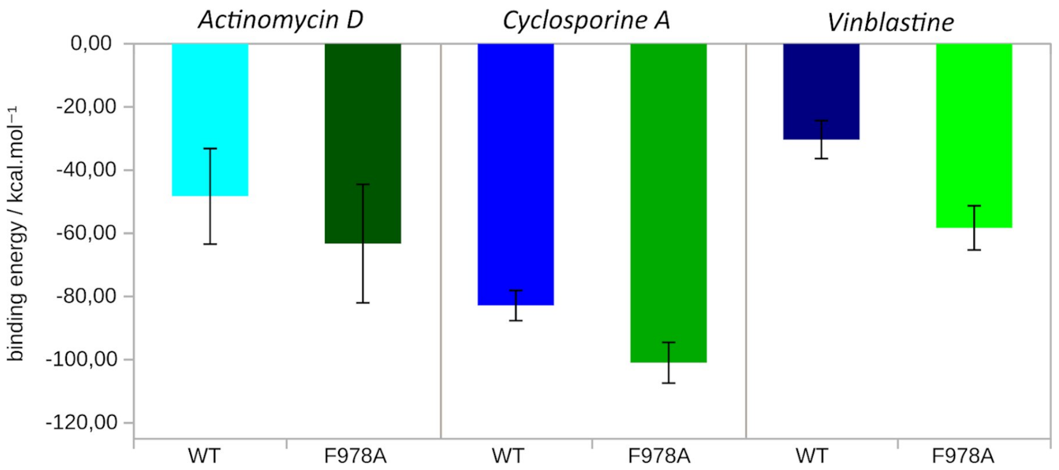


**Figure S51:** Relative Free energies of binding ($\Delta G_{bind}$) for actinomycin D, cyclosporine A and vinblastine in the WT and F978A variant.

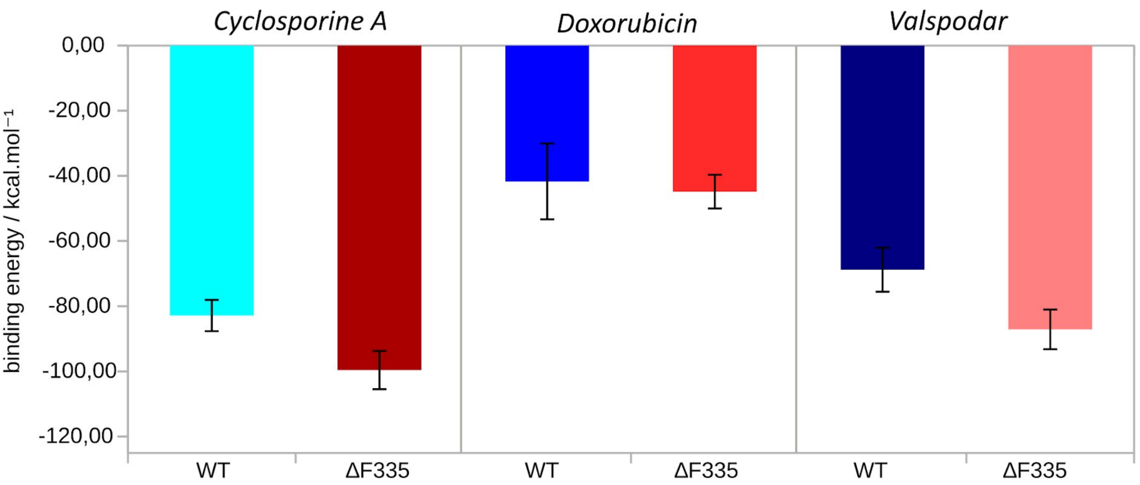


**Figure S52:** Relative Free energies of binding ($\Delta G_{bind}$) for cyclosporine A, doxorubicin and valspodar in the WT and ΔF335 variant.

**Table S25.** Variation in the contact frequencies and hydrogen bond (HB) parameters for colchicine (G185V) and vinblastine (G185V and F978A), both located at the H-site.

| Residues | | WT_COL | G185V_COL | | WT_VIN | G185V_VIN | | F978A_VIN | |
|---|---|---|---|---|---|---|---|---|---|
| | | Frequency | Frequency | Variation % | Frequency | Frequency | Variation % | Frequency | Variation % |
| 189 | LYS | 0,106931 | 0,683168 | 538,89 | 0,471287 | 0,332673 | -29,41 | 0,871287 | 84,87 |
| 192 | MET | | | | | 0,609901 | NEW | 0,263938 | NEW |
| 195 | GLN | | | | | 0,592079 | NEW | | |
| 196 | SER | | | | | 0,655446 | NEW | | |
| 344 | SER | | | | | 0,710891 | NEW | | |
| 347 | GLN | | | | | | | 0,648342 | NEW |
| 832 | ARG | | | | 0,522772 | | LOSS | 0,131747 | -74,80 |
| 938 | PHE | | | | | | | 0,824639 | NEW |
| 941 | THR | | | | | | | 0,726674 | NEW |
| 942 | PHE | 0,641584 | 0,269307 | -58,02 | | 0,847525 | NEW | 0,947510 | NEW |
| 945 | THR | 0,758416 | | LOSS | | | | | |
| 946 | GLN | | 0,825742 | NEW | | 0,570297 | NEW | 0,730289 | NEW |

| | | WT_COL | G185V_COL | Variation % | WT_VIN | G185V_VIN | Variation % | F978A_VIN | Variation % |
|---|---|---|---|---|---|---|---|---|---|
| HB | life | 1473,240 | 1063,813 | -27,79 | 383,12 | 1353,856 | 253,38 | 1177,059 | 207,23 |
| | <N> | 1,706 | 2,729 | 59,93 | 1,32 | 0,604 | -54,23 | 1,422 | 7,71 |
| | ΔG | -22,713 | -21,872 | -3,70 | -19,55 | -21,846 | 11,72 | -22,369 | 14,39 |

The variation percentage is estimated based on the mean frequencies of five replica systems for the *holo* P-gp systems. Green-shaded cells, no contact frequencies calculated for the specified residue.

**Table S26.** Variation in the contact frequencies and hydrogen bond (HB) parameters for actinomycin D (G830V and F978A) and doxorubicin (G830V and ΔF335), both located at the R-site.

| Residues | | WT_ACT | G830V_ACT | | F978A_ACT | | WT_DOX | G830V_DOX | | ΔF335_DOX | |
|---|---|---|---|---|---|---|---|---|---|---|---|
| | | Frequency | Frequency | Variation % | Frequency | Variation % | Frequency | Frequency | Variation % | Frequency | Variation % |
| 232 | TRP | 0,758416 | 0,829703 | 9,40 | 0,897030 | 18,28 | 0,500990 | 0,823779 | 64,43 | 0,650401 | 29,82 |
| 291 | LYS | 0,477228 | | LOSS | 0,304951 | -36,10 | | | | | |
| 296 | ASN | 0,164357 | | | 0,912871 | 455,42 | 0,875247 | 0,600358 | -31,41 | 1,000000 | 14,25 |
| 347 | GLN | 0,215842 | 0,576237 | 166,97 | | | | | | | |
| 770 | PHE | 0,306931 | 0,643564 | 109,68 | 0,514851 | 67,74 | | | | | |
| 773 | GLN | 0,209901 | 0,463366 | 120,75 | 0,619802 | 195,28 | 0,310891 | 0,905019 | 191,10 | 0,659088 | 112,00 |
| 777 | PHE | | | | | | 0,750495 | 0,703243 | -6,30 | 0,185971 | -75,22 |
| 778 | GLY | | | | | | 0,403960 | 0,561318 | 38,95 | | LOSS |
| 831 | SER | 0,431683 | 0,960396 | 122,48 | 0,568317 | 31,65 | | | | | |
| 838 | GLN | 0,253465 | 0,895049 | 253,12 | 0,400000 | 57,81 | 0,546534 | 0,548412 | 0,34 | 0,715486 | 30,91 |

| | | WT_ACT | G830V_ACT | Variation % | F978A_ACT | Variation % | WT_DOX | G830V_DOX | Variation % | ΔF335_DOX | Variation % |
|---|---|---|---|---|---|---|---|---|---|---|---|
| HB | life | 1062,751 | 525,676 | -50,54 | 406,804 | -61,72 | 397,677 | 561,385 | 41,17 | 1177,059 | 195,98 |
| | <N> | 2,638 | 2,443 | -7,36 | 2,897 | 9,83 | 0,896 | 0,774 | -13,64 | 1,422 | 58,68 |
| | ΔG | -21,956 | -19,878 | -9,47 | -19,694 | -10,30 | -19,649 | -20,086 | 2,22 | -22,369 | 13,85 |

The variation percentage is estimated based on the mean frequencies of five replica systems for the *holo* P-gp systems. Green-shaded cells, no contact frequencies calculated for the specified residue.

**Table S27.** Variation in the contact frequencies and hydrogen bond (HB) parameters for cyclosporine A (F978A and ΔF335 ) and valspodar (ΔF335), both located at the M-site.

| | | WT_CYC | F978A_CYC | | ΔF335_CYC | | WT_VLS | ΔF335_VLS | |
|---|---|---|---|---|---|---|---|---|---|
| Residues | | Frequency | Frequency | Variation % | Frequency | Variation % | Frequency | Frequency | Variation % |
| 303 | PHE | 0,522772 | 0,615408 | 17,72 | 0,691089 | 32,20 | 0,841584 | 0,388119 | -53,88 |
| 307 | TYR | 0,813861 | 0,765076 | -5,99 | 0,514852 | -36,74 | | | |
| 310 | TYR | 0,883168 | 0,662578 | -24,98 | 0,368317 | -58,30 | | | |
| 343 | PHE | 0,370297 | 0,678006 | 83,10 | 0,667327 | 80,21 | 0,419802 | 0,803960 | 91,51 |
| 347 | GLN | 0,819802 | 0,566839 | -30,86 | 0,445544 | -45,65 | 0,809901 | 0,536634 | -33,74 |
| 721 | ASN | | 0,604827 | NEW | | | | | |
| 725 | GLN | 0,164356 | 0,792936 | 382,45 | 0,663366 | 303,61 | 0,720792 | 0,283168 | -60,71 |
| 759 | PHE | | | | | | 0,605940 | | LOSS |
| 766 | SER | | 0,681897 | NEW | | | | | |
| 770 | PHE | 0,295050 | 0,460815 | 56,18 | | | | | |
| 946 | GLN | 0,174258 | | | 0,516832 | 196,59 | | | |
| 983 | PHE | | | | | | 0,576238 | 0,368317 | -36,08 |
| 990 | GLN | | | | | | 0,097030 | 0,641584 | 561,22 |

| | | WT_CYC | F978A_CYC | Variation % | ΔF335_CYC | Variation % | WT_VLS | ΔF335_VLS | Variation % |
|---|---|---|---|---|---|---|---|---|---|
| HB | life | 624,098 | 602,512 | -3,46 | 644,905 | 3,33 | 887,101 | 770,965 | -13,09 |
| | <N> | 2,879 | 2,822 | -1,99 | 2,255 | -21,69 | 1,500 | 1,774 | 18,29 |
| | ΔG | -20,614 | -20,587 | -0,13 | -20,811 | 0,95 | -21,478 | -21,510 | 0,15 |

*The variation percentage is estimated based on the mean frequencies of five replica systems for the holo* P-gp systems. Green-shaded cells, no contact frequencies calculated for the specified residue.

**Table S28**: Protein-ligand contact efficiency ratio for the WT and P-gp variants.

| | WT | | | G185V_H | | |
|---|---|---|---|---|---|---|
| Ligand | mean Total | Mean (>0,5) | Ratio | mean Total | Mean (>0,5) | Ratio |
| COL_H | 8,80 | 3,80 | 0,43 | 9,80 | 4,20 | 0,43 |
| VIN_H | 13,20 | 4,00 | 0,30 | 8,00 | 5,60 | 0,70 |

| | WT | | | G830V_R | | |
|---|---|---|---|---|---|---|
| Ligand | mean Total | Mean (>0,5) | Ratio | mean Total | Mean (>0,5) | Ratio |
| ACT_R | 19,60 | 4,00 | 0,20 | 18,20 | 6,20 | 0,34 |
| DOX_R | 19,80 | 5,80 | 0,29 | 20,60 | 6,80 | 0,33 |

| | WT | | | F978A_M | | |
|---|---|---|---|---|---|---|
| Ligand | mean Total | Mean (>0,5) | Ratio | mean Total | Mean (>0,5) | Ratio |
| ACT_R | 19,60 | 4,00 | 0,20 | 19,80 | 5,20 | 0,26 |
| CYC_M | 27,60 | 3,80 | 0,14 | 26,60 | 7,60 | 0,29 |
| VIN_H | 13,20 | 4,00 | 0,30 | 16,60 | 7,00 | 0,42 |

| | WT | | | ΔF335_M | | |
|---|---|---|---|---|---|---|
| Ligand | mean Total | Mean (>0,5) | Ratio | mean Total | Mean (>0,5) | Ratio |
| CYC_M | 27,60 | 3,80 | 0,14 | 26,60 | 6,60 | 0,25 |
| DOX_R | 19,80 | 5,80 | 0,29 | 19,20 | 5,20 | 0,27 |
| VLS_M | 22,60 | 5,40 | 0,24 | 25,20 | 6,40 | 0,25 |

The variation percentage is estimated based on the mean frequencies of five replica systems for the *holo* P-gp systems.

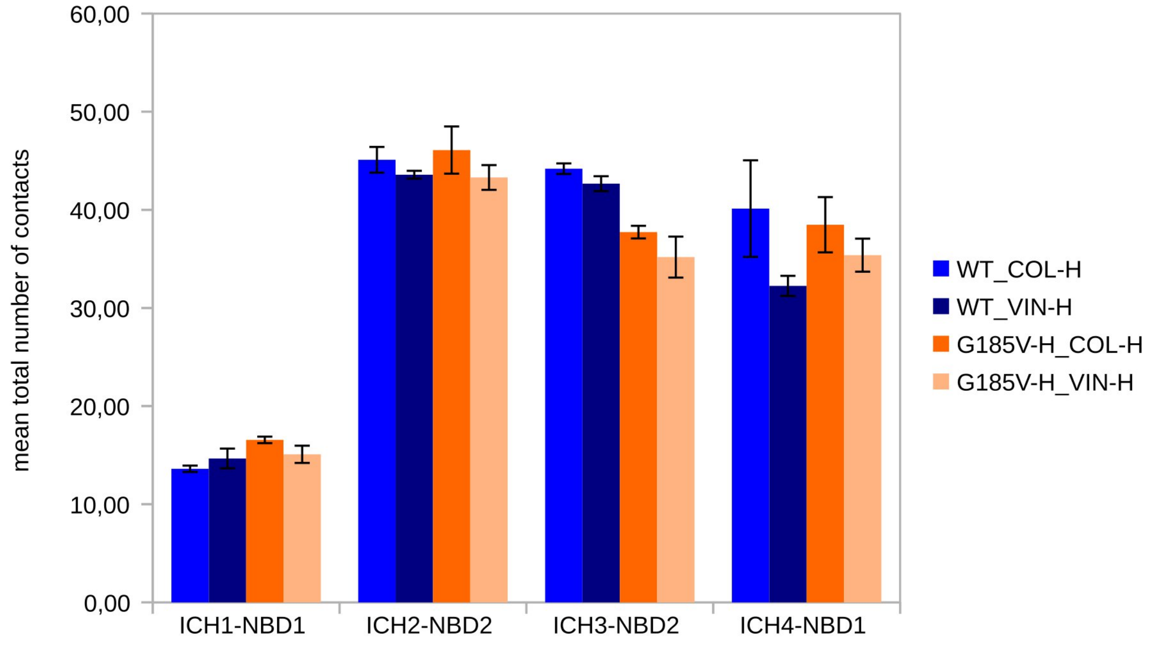


**Figure S53:** Comparison of the total number of contacts at each ICH-NBD interface between the WT and G185V variant (*holo* systems).

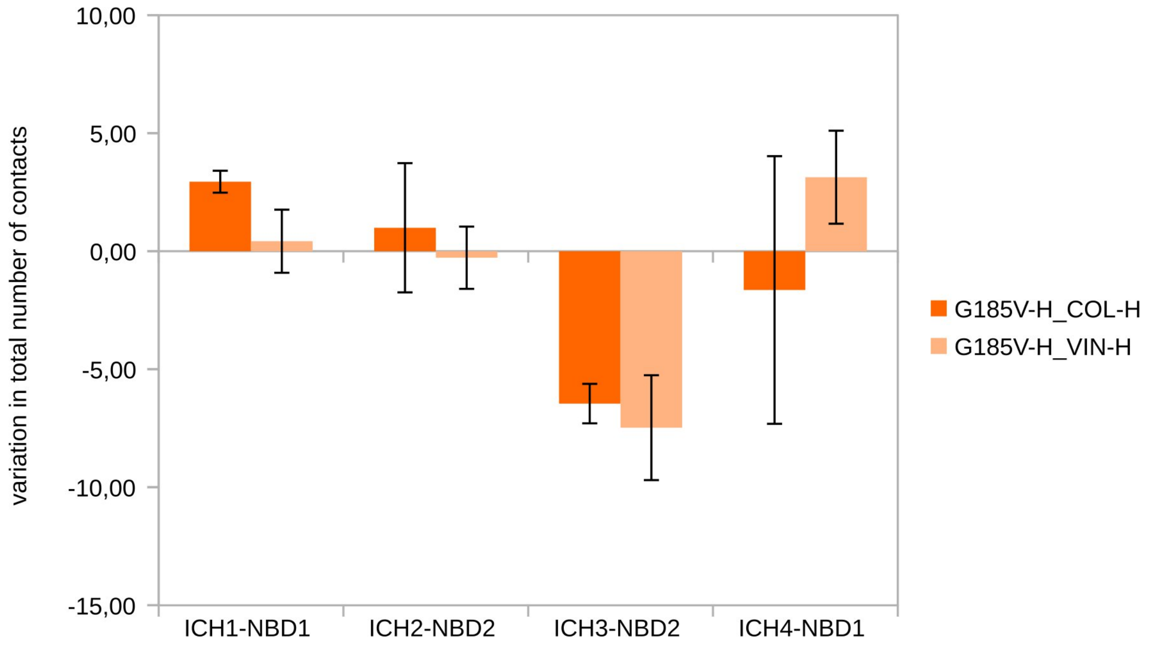


**Figure S54:** Variation in the total number of contacts at each ICH-NBD interface for the *holo* G185V variant (difference from the *holo* WT).

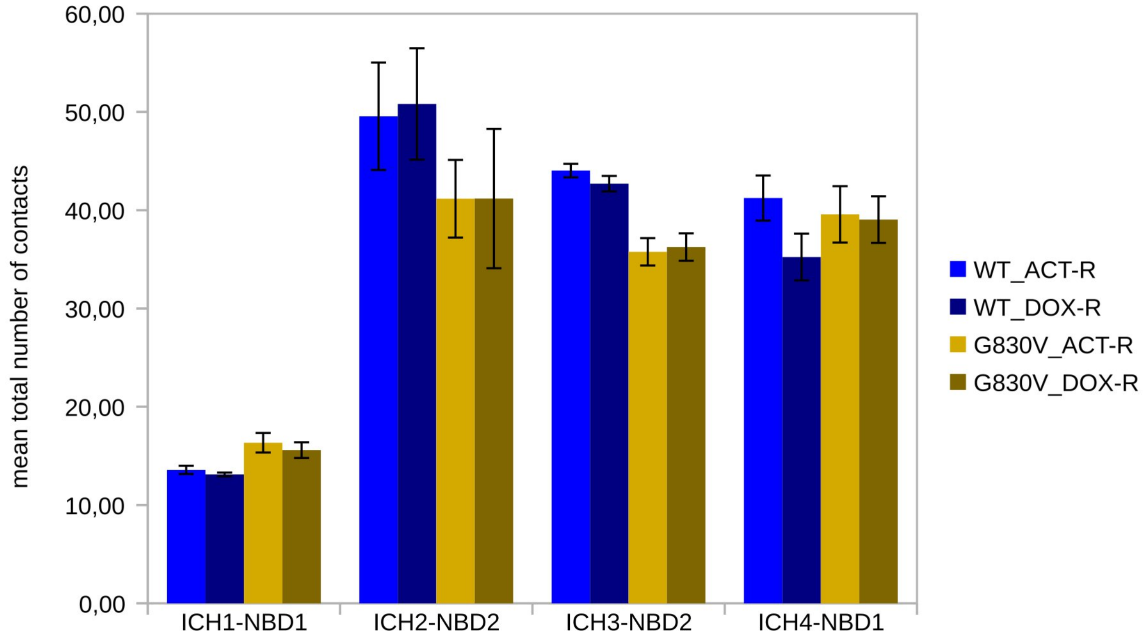


**Figure S55:** Comparison of the total number of contacts at each ICH-NBD interface between the WT and G830V variant (*holo* systems).

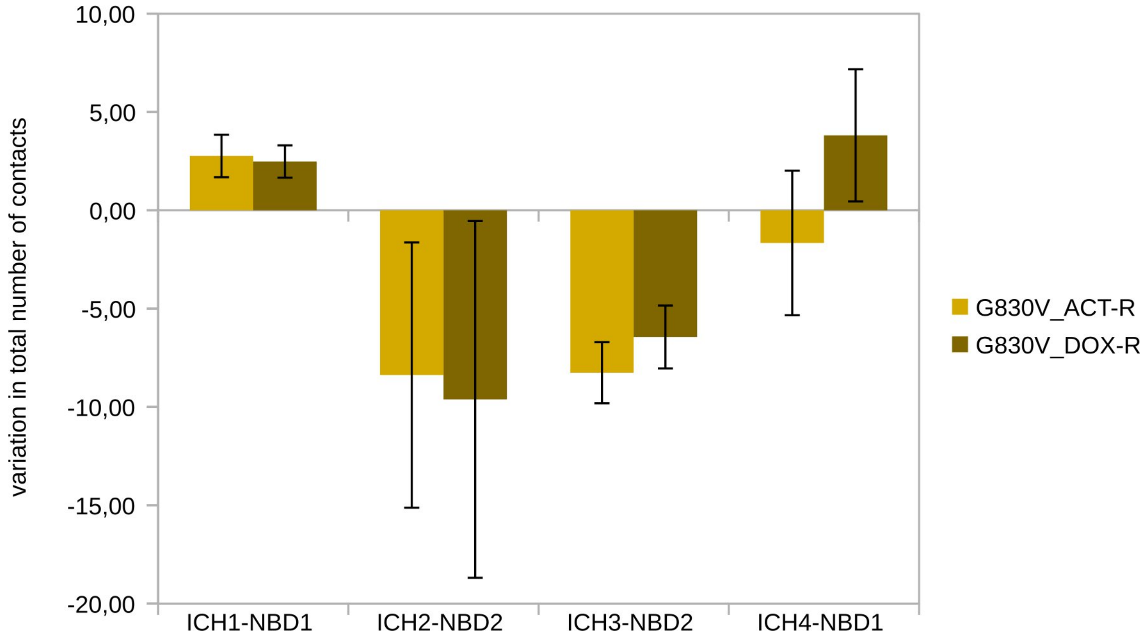


**Figure S56:** Variation in the total number of contacts at each ICH-NBD interface for the *holo* G830V variant (difference from the *holo* WT).

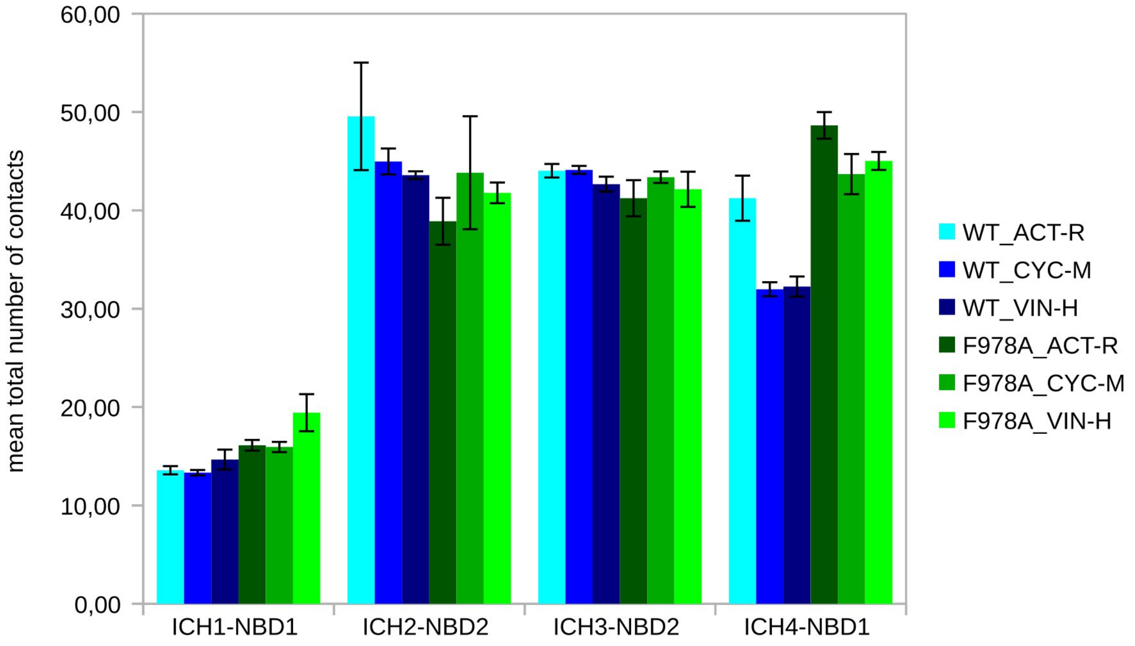


**Figure S57:** Comparison of the total number of contacts at each ICH-NBD interface between the WT and F978A variant (*holo* systems).

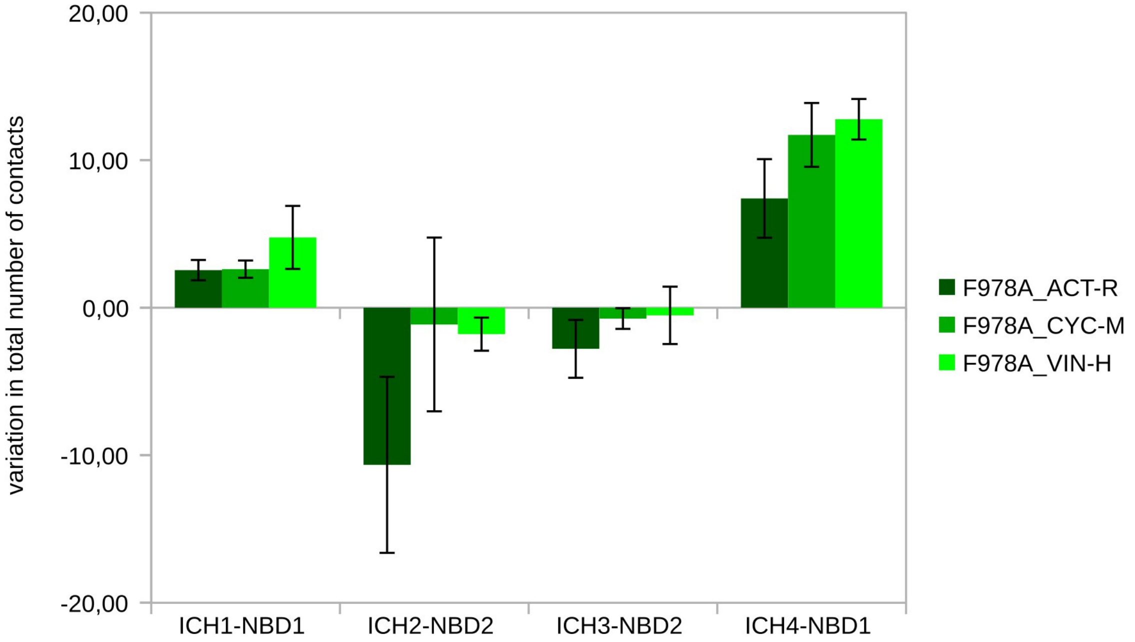


**Figure S58:** Variation in the total number of contacts at each ICH-NBD interface for the *holo* F978A variant (difference from the *holo* WT).

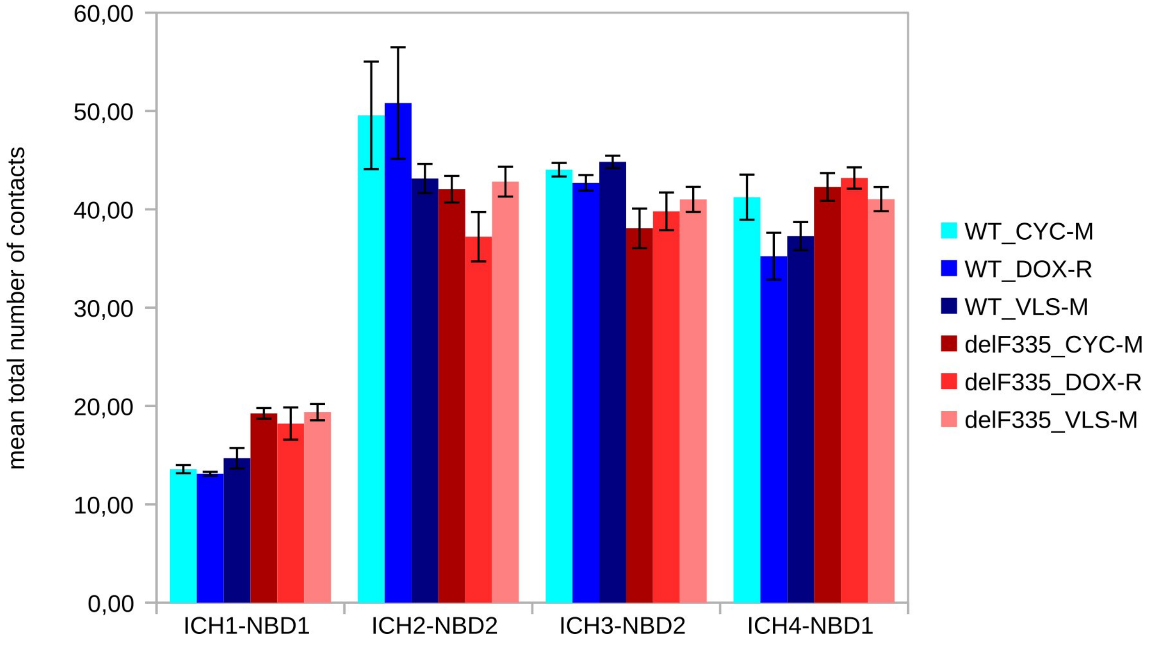


**Figure S59:** Comparison of the total number of contacts at each ICH-NBD interface between the WT and ΔF335 variant (*holo* systems).

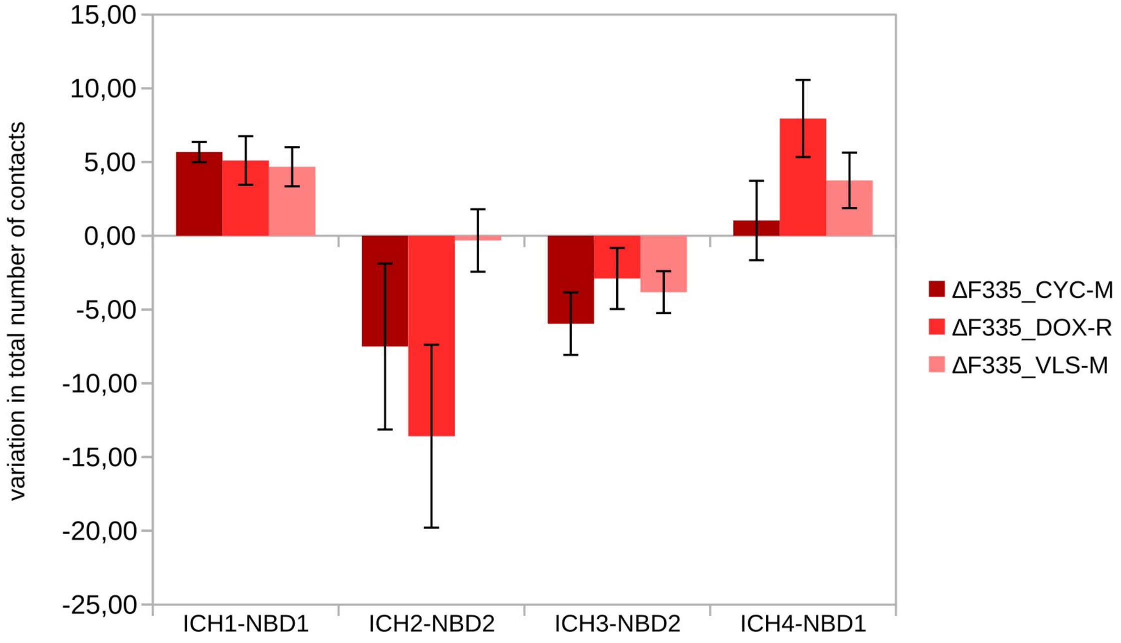


**Figure S60:** Variation in the total number of contacts at each ICH-NBD interface for the *holo* ΔF335 variant (difference from the *holo* WT).

**Table S29.** Variation in the contact frequencies and hydrogen bond (HB) parameters between the ICH-NBD residues for the WT and G185V variants (*holo* systems).

| TMD-NBD interface | | | | WT_COL | G185V_COL | | WT_VIN | G185V_VIN | |
|---|---|---|---|---|---|---|---|---|---|
| ICH1 | | NBD1 | | Frequency | Frequency | Variation % | Frequency | Frequency | Variation % |
| 160 | ILE | 443 | LEU | 0,188119 | 0,748515 | 297,89 | 0,356436 | 0,502970 | 41,11 |
| 164 | ASP | 404 | ARG | 0,946535 | 0,982178 | 3,77 | 0,970297 | 0,859406 | -11,43 |
| 164 | ASP | 405 | LYS | 0,629505 | 0,984158 | 56,34 | 0,960396 | 0,786139 | -18,14 |

| Hydrogen Bonds | | WT_COL | G185V_COL | Variation % | WT_VIN | G185V_VIN | Variation % |
|---|---|---|---|---|---|---|---|
| ICH1-NBD1 | life | 570,612 | 709,366 | 24,32 | 426,906 | 369,544 | -13,44 |
| | <N> | 1,911 | 2,442 | 27,77 | 2,188 | 1,976 | -9,69 |
| | DG | -20,376 | -20,730 | 1,74 | -19,734 | -19,247 | -2,47 |

| TMD-NBD interface | | | | WT_COL | G185V_COL | | WT_VIN | G185V_VIN | |
|---|---|---|---|---|---|---|---|---|---|
| ICH2 | | NBD2 | | Frequency | Frequency | Variation % | Frequency | Frequency | Variation % |
| 262 | ARG | 1086 | PHE | 0,425743 | 0,576238 | 35,35 | 0,502970 | 0,568317 | 12,99 |
| 263 | THR | 1117 | SER | 0,635643 | 0,603168 | -5,11 | 0,649505 | 0,611881 | -5,79 |
| 263 | THR | 1118 | GLN | 0,035644 | 0,423762 | 1088,89 | 0,031683 | 0,512871 | 1518,75 |
| 263 | THR | 1200 | ASP | 0,198020 | 0,578218 | 192,00 | 0,031683 | 0,673267 | 2025,00 |
| 265 | ILE | 1110 | ARG | 0,758416 | 0,657426 | -13,32 | 0,639604 | 0,623762 | -2,48 |
| 266 | ALA | 1086 | PHE | 0,596039 | 0,483169 | -18,94 | 0,596039 | 0,522772 | -12,29 |
| 267 | PHE | 1110 | ARG | 0,504911 | 0,332674 | -34,11 | 0,471287 | 0,526733 | 11,76 |
| 267 | PHE | 1115 | ILE | 0,936634 | 0,908911 | -2,96 | 0,897030 | 0,920792 | 2,65 |

| Hydrogen Bonds | | WT_COL | G185V_COL | Variation % | WT_VIN | G185V_VIN | Variation % |
|---|---|---|---|---|---|---|---|
| ICH2-NBD2 | life | 452,519 | 393,494 | -13,04 | 335,881 | 940,684 | 180,06 |
| | <N> | 1,396 | 1,861 | 33,30 | 1,095 | 1,600 | 46,13 |
| | DG | -19,839 | -19,057 | -3,94 | -18,981 | -20,133 | 6,07 |

| TMD-NBD interface | | | | WT_COL | G185V_COL | | WT_VIN | G185V_VIN | |
|---|---|---|---|---|---|---|---|---|---|
| ICH3 | | NBD2 | | Frequency | Frequency | Variation % | Frequency | Frequency | Variation % |
| 800 | ASP | 1086 | PHE | 0,736633 | 0,826733 | 12,23 | 0,699010 | 0,782178 | 11,90 |
| 800 | ASP | 1087 | TYR | 0,994059 | 0,995050 | 0,10 | 0,990099 | 0,995050 | 0,50 |
| 801 | VAL | 1086 | PHE | 0,902970 | 0,883663 | -2,14 | 0,914851 | 0,873762 | -4,49 |
| 801 | VAL | 1087 | TYR | 0,469307 | 0,512376 | 9,18 | 0,485149 | 0,502475 | 3,57 |
| 802 | SER | 1087 | TYR | 0,687128 | 0,646039 | -5,98 | 0,693069 | 0,695544 | 0,36 |
| 805 | ASP | 1044 | TYR | 0,912871 | 0,908416 | -0,49 | 0,922772 | 0,888614 | -3,70 |
| 805 | ASP | 1046 | THR | 1,000000 | 0,878713 | -12,13 | 1,000000 | 0,626238 | -37,38 |

| Hydrogen Bonds | | WT_COL | G185V_COL | Variation % | WT_VIN | G185V_VIN | Variation % |
|---|---|---|---|---|---|---|---|
| ICH3-NBD2 | life | 981,591 | 690,718 | -29,63 | 702,618 | 339,698 | -51,65 |
| | <N> | 2,814 | 1,790 | -36,37 | 2,871 | 1,956 | 0,96 |
| | DG | -20,899 | -20,815 | -0,40 | -20,761 | -19,098 | -20,10 |

| TMD-NBD interface | | | | WT_COL | G185V_COL | | WT_VIN | G185V_VIN | |
|---|---|---|---|---|---|---|---|---|---|
| ICH4 | | NBD1 | | Frequency | Frequency | Variation % | Frequency | Frequency | Variation % |
| 905 | ARG | 441 | GLN | 1,000000 | 1,000000 | 0,00 | 1,000000 | 1,000000 | 0,00 |
| 908 | VAL | 467 | ARG | 0,736633 | 0,697029 | -5,38 | 0,609901 | 0,631683 | 3,57 |
| 909 | SER | 441 | GLN | 0,998020 | 0,839604 | -15,87 | 1,000000 | 0,887129 | -11,29 |
| 911 | THR | 467 | ARG | 0,724752 | 0,000000 | Lost | 0,459406 | 0,057426 | -87,50 |
| 912 | GLN | 464 | ARG | 0,623762 | 0,318812 | -48,89 | 0,435644 | 0,366337 | -15,91 |
| 912 | GLN | 467 | ARG | 0,150495 | 0,572277 | 280,26 | 0,180198 | 0,394059 | 118,68 |

| Hydrogen Bonds | | WT_COL | G185V_COL | Variation % | WT_VIN | G185V_VIN | Variation % |
|---|---|---|---|---|---|---|---|
| ICH4-NBD1 | life | 475,834 | 365,229 | -23,24 | 454,402 | 284,478 | -37,40 |
| | <N> | 3,063 | 2,673 | -12,73 | 2,556 | 1,991 | -22,14 |
| | DG | -19,742 | -18,657 | -5,50 | -19,648 | -18,789 | -4,37 |

The variation percentage is estimated based on the mean frequencies of five replica systems for the WT and G185V variant (*holo* systems).

**Table S30.** Variation in the contact frequencies and hydrogen bond (HB) parameters between the ICH-NBD residues for the WT and G830V variants (*holo* systems).

| TMD-NBD interface | | | | WT_ACT | G830V_ACT | | WT_DOX | G830V_DOX | |
|---|---|---|---|---|---|---|---|---|---|
| ICH1 | | NBD1 | | Frequency | Frequency | Variation % | Frequency | Frequency | Variation % |
| 164 | ASP | 404 | ARG | 0,984139 | 0,986149 | 0,20 | 0,986119 | 0,992060 | 0,60 |
| 164 | ASP | 405 | LYS | 0,991074 | 0,841535 | -15,09 | 0,940535 | 0,828688 | -11,89 |

| Hydrogen Bonds | | WT_ACT | G830V_ACT | Variation % | WT_DOX | G830V_DOX | Variation % |
|---|---|---|---|---|---|---|---|
| ICH1-NBD1 | life | 779,200 | 677,000 | -13,12 | 724,200 | 756,000 | 4,39 |
| | <N> | 2,035 | 2,065 | 1,50 | 2,020 | 2,009 | -0,55 |
| | DG | -21,072 | -20,895 | -0,84 | -21,022 | -21,339 | 1,50 |

| TMD-NBD interface | | | | WT_ACT | G830V_ACT | | WT_DOX | G830V_DOX | |
|---|---|---|---|---|---|---|---|---|---|
| ICH2 | | NBD2 | | Frequency | Frequency | Variation % | Frequency | Frequency | Variation % |
| 262 | ARG | 1077 | SER | 0,463366 | 0,269307 | -41,88 | 0,000000 | 0,000000 | 0,00 |
| 262 | ARG | 1081 | GLN | 0,475248 | 0,401980 | -15,42 | 0,449505 | 0,334500 | -25,58 |
| 262 | ARG | 1086 | PHE | 0,457426 | 0,520792 | 13,85 | 0,481188 | 0,448378 | -6,82 |
| 263 | THR | 1117 | SER | 0,540594 | 0,316832 | -41,39 | 0,744554 | 0,374684 | -49,68 |
| 265 | ILE | 1110 | ARG | 0,695049 | 0,740594 | 6,55 | 0,716831 | 0,860754 | 20,08 |
| 266 | ALA | 1086 | PHE | 0,623762 | 0,459386 | -26,35 | 0,532673 | 0,462209 | -13,23 |
| 266 | ALA | 1110 | ARG | 0,421783 | 0,548515 | 30,05 | 0,459406 | 0,415108 | -9,64 |
| 267 | PHE | 1110 | ARG | 0,461386 | 0,429703 | -6,87 | 0,477228 | 0,327416 | -31,39 |
| 267 | PHE | 1115 | ILE | 0,916832 | 0,827723 | -9,72 | 0,934653 | 0,884585 | -5,36 |

| Hydrogen Bonds | | WT_ACT | G830V_ACT | Variation % | WT_DOX | G830V_DOX | Variation % |
|---|---|---|---|---|---|---|---|
| ICH2-NBD2 | life | 301,800 | 1782,600 | 490,66 | 501,600 | 496,800 | -0,96 |
| | <N> | 1,142 | 1,104 | -3,38 | 1,309 | 0,980 | -25,19 |
| | DG | -18,851 | -22,547 | 19,61 | -20,198 | -19,927 | -1,34 |

| TMD-NBD interface | | | | WT_ACT | G830V_ACT | | WT_DOX | G830V_DOX | |
|---|---|---|---|---|---|---|---|---|---|
| ICH3 | | NBD2 | | Frequency | Frequency | Variation % | Frequency | Frequency | Variation % |
| 800 | ASP | 1086 | PHE | 0,754455 | 0,780198 | 3,41 | 0,718812 | 0,794452 | 10,52 |
| 800 | ASP | 1087 | TYR | 1,000000 | 0,998020 | -0,20 | 0,968317 | 0,994059 | 2,66 |
| 801 | VAL | 1086 | PHE | 0,928713 | 0,853465 | -8,10 | 0,891089 | 0,885029 | -0,68 |
| 801 | VAL | 1087 | TYR | 0,526733 | 0,091089 | -82,71 | 0,510891 | 0,032127 | -93,71 |
| 802 | SER | 1087 | TYR | 0,736633 | 0,299010 | -59,41 | 0,655445 | 0,279891 | -57,30 |
| 805 | ASP | 1044 | TYR | 0,904950 | 0,469307 | -48,14 | 0,930693 | 0,504916 | -45,75 |
| 805 | ASP | 1046 | THR | 0,998020 | 0,998020 | 0,00 | 0,998020 | 0,803960 | -19,44 |

| Hydrogen Bonds | | WT_ACT | G830V_ACT | Variation % | WT_DOX | G830V_DOX | Variation % |
|---|---|---|---|---|---|---|---|
| ICH3-NBD2 | life | 858,200 | 773,800 | -9,83 | 565,400 | 623,800 | 10,33 |
| | <N> | 2,849 | 2,979 | 4,56 | 2,686 | 2,997 | 11,59 |
| | DG | -21,282 | -21,316 | 0,16 | -20,742 | -20,789 | 0,22 |

| TMD-NBD interface | | | | WT_ACT | G830V_ACT | | WT_DOX | G830V_DOX | |
|---|---|---|---|---|---|---|---|---|---|
| ICH4 | | NBD1 | | Frequency | Frequency | Variation % | Frequency | Frequency | Variation % |
| 905 | ARG | 438 | GLN | 0,451485 | 0,146535 | -67,54 | 0,000000 | 0,000000 | 0,00 |
| 905 | ARG | 441 | GLN | 1,000000 | 0,998020 | -0,20 | 0,998020 | 0,996040 | -0,20 |
| 908 | VAL | 467 | ARG | 0,566337 | 0,720811 | 27,28 | 0,560396 | 0,708928 | 26,50 |
| 909 | SER | 441 | GLN | 0,992079 | 0,912871 | -7,98 | 0,980198 | 0,868163 | -11,43 |
| 910 | LEU | 467 | ARG | 0,000000 | 0,936634 | NEW | 0,000000 | 0,828883 | NEW |
| 911 | THR | 467 | ARG | 0,617830 | 0,376238 | -39,10 | 0,000000 | 0,000000 | 0,00 |
| 912 | GLN | 464 | ARG | 0,643564 | 0,467327 | -27,38 | 0,000000 | 0,000000 | 0,00 |

| Hydrogen Bonds | | WT_ACT | G830V_ACT | Variation % | WT_DOX | G830V_DOX | Variation % |
|---|---|---|---|---|---|---|---|
| ICH4-NBD1 | life | 621,200 | 441,600 | -28,91 | 365,200 | 705,400 | 93,15 |
| | <N> | 2,828 | 2,495 | -11,78 | 2,544 | 2,368 | -6,90 |
| | DG | -20,458 | -19,820 | -3,12 | -19,324 | -20,750 | 7,38 |

The variation percentage is estimated based on the mean frequencies of five replica systems for the WT and G830V variant (*holo* systems).

**Table S31.** Variation in the contact frequencies and hydrogen bond (HB) parameters between the ICH-NBD residues for the WT and F978A variants (*holo* systems).

| TMD-NBD interface | | | | WT_ACT | F978A_ACT | | WT_CYC | F978A_CYC | | WT_VIN | F978A_VIN | |
|---|---|---|---|---|---|---|---|---|---|---|---|---|
| ICH1 | | NBD1 | | Frequency | Frequency | Variation % | Frequency | Frequency | Variation % | Frequency | Frequency | Variation % |
| 160 | ILE | 443 | LEU | 0,000000 | 0,000000 | 0,00 | 0,140594 | 0,485149 | 245,07 | 0,356436 | 0,661386 | 85,56 |
| 164 | ASP | 404 | ARG | 0,984139 | 0,936614 | -4,83 | 0,974257 | 0,964356 | -1,02 | 0,970297 | 0,980198 | 1,02 |
| 164 | ASP | 405 | LYS | 0,991074 | 0,776238 | -21,68 | 0,990099 | 0,871287 | -12,00 | 0,960396 | 0,778218 | -18,97 |

| Hydrogen Bonds | | WT_ACT | F978A_ACT | Variation % | WT_CYC | F978A_CYC | Variation % | WT_VIN | F978A_VIN | Variation % |
|---|---|---|---|---|---|---|---|---|---|---|
| ICH1-NBD1 | life | 779,200 | 442,800 | -43,17 | 484,600 | 520,400 | 7,39 | 426,9058 | 392,800 | -7,99 |
| | <N> | 2,035 | 2,038 | 0,15 | 1,929 | 2,131 | 10,46 | 2,1882 | 2,442 | 11,59 |
| | DG | -21,072 | -19,931 | -5,41 | -20,150 | -20,238 | 0,44 | -19,7344 | -19,572 | -0,82 |

| TMD-NBD interface | | | | WT_ACT | F978A_ACT | | WT_CYC | F978A_CYC | | WT_VIN | F978A_VIN | |
|---|---|---|---|---|---|---|---|---|---|---|---|---|
| ICH2 | | NBD2 | | Frequency | Frequency | Variation % | Frequency | Frequency | Variation % | Frequency | Frequency | Variation % |
| 262 | ARG | 1044 | TYR | 0,000000 | 0,000000 | 0,000000 | 0,000000 | 0,000000 | 0,000000 | 0,5148514 | 0,403961 | -21,54 |
| 262 | ARG | 1077 | SER | 0,463366 | 0,435643 | -5,98 | 0,447525 | 0,560396 | 25,22 | 0,3841586 | 0,463367 | 20,62 |
| 262 | ARG | 1081 | GLN | 0,475248 | 0,495050 | 4,17 | 0,407921 | 0,594099 | 45,64 | 0,3960398 | 0,550495 | 39,00 |
| 262 | ARG | 1086 | PHE | 0,457426 | 0,465347 | 1,73 | 0,514851 | 0,439604 | -14,62 | 0,5029704 | 0,550495 | 9,45 |
| 263 | THR | 1117 | SER | 0,540594 | 0,562376 | 4,03 | 0,740594 | 0,633663 | -14,44 | 0,6495048 | 0,526733 | -18,90 |
| 265 | ILE | 1110 | ARG | 0,695049 | 0,396040 | -43,02 | 0,738614 | 0,467327 | -36,73 | 0,6396038 | 0,370297 | -42,11 |
| 266 | ALA | 1086 | PHE | 0,623762 | 0,487129 | -21,90 | 0,600000 | 0,510891 | -14,85 | 0,5960394 | 0,459406 | -22,92 |
| 266 | ALA | 1110 | ARG | 0,000000 | 0,000000 | 0,000000 | 0,000000 | 0,000000 | 0,000000 | 0,4534656 | 0,437624 | -3,49 |
| 267 | PHE | 1110 | ARG | 0,461386 | 0,336634 | -27,04 | 0,560396 | 0,457426 | -18,37 | 0,4712874 | 0,310891 | -34,03 |
| 267 | PHE | 1115 | ILE | 0,9168316 | 0,8534652 | -6,91 | 0,944574 | 0,851485 | -9,86 | 0,8970296 | 0,853465 | -4,86 |

| Hydrogen Bonds | | WT_ACT | F978A_ACT | Variation % | WT_CYC | F978A_CYC | Variation % | WT_VIN | F978A_VIN | |
|---|---|---|---|---|---|---|---|---|---|---|
| ICH2-NBD2 | life | 301,800 | 325,000 | 7,69 | 497,600 | 398,800 | -19,86 | 335,881 | 317,8 | -5,38 |
| | <N> | 1,142 | 1,053 | -7,79 | 1,367 | 1,774 | 29,83 | 1,0948 | 1,285 | 17,37 |
| | DG | -18,851 | -19,120 | 1,43 | -20,078 | -19,578 | -2,49 | -18,9814 | -18,648 | -1,76 |

| TMD-NBD interface | | | | WT_ACT | F978A_ACT | | WT_CYC | F978A_CYC | | WT_VIN | F978A_VIN | |
|---|---|---|---|---|---|---|---|---|---|---|---|---|
| ICH3 | | NBD2 | | Frequency | Frequency | Variation % | Frequency | Frequency | Variation % | Frequency | Frequency | Variation % |
| 800 | ASP | 1086 | PHE | 0,754455 | 0,720792 | -4,46 | 0,734653 | 0,726732 | -1,08 | 0,699010 | 0,768317 | 9,92 |
| 800 | ASP | 1087 | TYR | 1,000000 | 0,996040 | -0,40 | 0,998020 | 1,000000 | 0,20 | 0,990099 | 0,984158 | -0,60 |
| 801 | VAL | 1086 | PHE | 0,928713 | 0,893069 | -3,84 | 0,899010 | 0,893069 | -0,66 | 0,914851 | 0,869247 | -4,98 |
| 801 | VAL | 1087 | TYR | 0,526733 | 0,433663 | -17,67 | 0,489109 | 0,508911 | 4,05 | 0,485149 | 0,516831 | 6,53 |
| 802 | SER | 1087 | TYR | 0,736633 | 0,516831 | -29,84 | 0,685148 | 0,643564 | -6,07 | 0,693069 | 0,568317 | -18,00 |
| 805 | ASP | 1044 | TYR | 0,904950 | 0,851485 | -5,91 | 0,936634 | 0,904950 | -3,38 | 0,922772 | 0,869307 | -5,79 |
| 805 | ASP | 1046 | THR | 0,998020 | 0,998020 | 0,00 | 0,998020 | 0,998020 | 0,00 | 1,000000 | 1,000000 | 0,00 |

| Hydrogen Bonds | | WT_ACT | F978A_ACT | Variation % | WT_CYC | F978A_CYC | Variation % | WT_VIN | F978A_VIN | |
|---|---|---|---|---|---|---|---|---|---|---|
| ICH3-NBD2 | life | 858,200 | 471,800 | -45,02 | 822,600 | 388,800 | -52,74 | 702,618 | 547,400 | -22,09 |
| | <N> | 2,849 | 2,709 | -4,93 | 2,841 | 2,568 | -9,61 | 2,871 | 2,509 | -12,61 |
| | DG | -21,282 | -19,803 | -6,95 | -21,464 | -19,488 | -9,21 | -20,761 | -19,792 | -4,67 |

| TMD-NBD interface | | | | WT_ACT | F978A_ACT | | WT_CYC | F978A_CYC | | WT_VIN | F978A_VIN | |
|---|---|---|---|---|---|---|---|---|---|---|---|---|
| ICH4 | | NBD1 | | Frequency | Frequency | Variation % | Frequency | Frequency | Variation % | Frequency | Frequency | Variation % |
| 905 | ARG | 401 | TYR | 0,174258 | 0,732655 | 320,44 | 0,217822 | 0,617822 | 183,64 | 0,220222 | 0,691089 | 213,81 |
| 905 | ARG | 438 | GLN | 0,451485 | 0,855426 | 89,47 | 0,215842 | 0,772277 | 257,80 | 0,245545 | 0,920792 | 275,00 |
| 905 | ARG | 441 | GLN | 1,000000 | 1,000000 | 0,00 | 0,964356 | 1,000000 | 3,70 | 1,000000 | 0,998020 | -0,20 |
| 908 | VAL | 467 | ARG | 0,566337 | 0,752475 | 32,87 | 0,568317 | 0,726732 | 27,87 | 0,609901 | 0,679208 | 11,36 |
| 909 | SER | 441 | GLN | 0,992079 | 1,000000 | 0,80 | 0,998020 | 0,996040 | -0,20 | 1,000000 | 0,994059 | -0,59 |
| 911 | THR | 467 | ARG | 0,617830 | 0,904950 | 46,47 | 0,352475 | 0,976238 | 176,97 | 0,459406 | 0,847525 | 84,48 |
| 912 | GLN | 464 | ARG | 0,643564 | 0,4752474 | -26,15 | 0,409901 | 0,400000 | -2,42 | 0,435644 | 0,398040 | -8,63 |

| Hydrogen Bonds | | WT_ACT | F978A_ACT | Variation % | WT_CYC | F978A_CYC | Variation % | WT_VIN | F978A_VIN | |
|---|---|---|---|---|---|---|---|---|---|---|
| ICH4-NBD1 | life | 621,200 | 446,400 | -28,14 | 476,000 | 347,800 | -26,93 | 454,402 | 324,200 | -28,65 |
| | <N> | 2,828 | 3,743 | 32,33 | 2,457 | 3,434 | 39,73 | 2,556 | 3,531 | 38,12 |
| | DG | -20,458 | -19,897 | -2,74 | -19,526 | -19,268 | -1,32 | -19,648 | -18,998 | -3,31 |

The variation percentage is estimated based on the mean frequencies of five replica systems for the WT and F978A variant (*holo* systems).

**Table S32.** Variation in the contact frequencies and hydrogen bond (HB) parameters between the ICH-NBD residues for the WT and ΔF335 variant (*holo* systems).

| TMD-NBD interface | | | | WT_CYC | ΔF335_CYC | | WT_VLS | ΔF335_VLS | | WT_DOX | ΔF335_DOX | |
|---|---|---|---|---|---|---|---|---|---|---|---|---|
| ICH1 | | NBD1 | | Frequency | Frequency | Variation % | Frequency | Frequency | Variation % | Frequency | Frequency | Variation % |
| 160 | ILE | 443 | LEU | 0,187183 | 0,613861 | 227,95 | 0,396040 | 0,568317 | 43,50 | 0,183518 | 0,513282 | 179,69 |
| 164 | ASP | 404 | ARG | 0,984139 | 0,984158 | 0,00 | 0,910891 | 0,976238 | 7,17 | 0,986119 | 0,980198 | -0,60 |
| 164 | ASP | 405 | LYS | 0,991074 | 0,000000 | LOST | 0,825743 | 0,000000 | LOST | 0,940535 | 0,000000 | LOST |

| Hydrogen Bonds | | WT_CYC | ΔF335_CYC | Variation % | WT_VLS | ΔF335_VLS | Variation % | WT_DOX | ΔF335_DOX | Variation % |
|---|---|---|---|---|---|---|---|---|---|---|
| ICH1-NBD1 | life | 779,200 | 217,800 | -72,05 | 427,400 | 227,200 | -46,84 | 724,200 | 440,200 | -39,22 |
| | <N> | 2,035 | 2,121 | 4,22 | 2,109 | 2,067 | -1,97 | 2,020 | 1,957 | -3,09 |
| | DG | -21,072 | -18,168 | -13,78 | -19,843 | -18,241 | -8,07 | -21,022 | -18,994 | -9,65 |

| TMD-NBD interface | | | | WT_CYC | ΔF335_CYC | | WT_VLS | ΔF335_VLS | | WT_DOX | ΔF335_DOX | |
|---|---|---|---|---|---|---|---|---|---|---|---|---|
| ICH2 | | NBD2 | | Frequency | Frequency | Variation % | Frequency | Frequency | Variation % | Frequency | Frequency | Variation % |
| 262 | ARG | 1077 | SER | 0,447525 | 0,536633 | 19,91 | 0,461386 | 0,536633 | 16,31 | 0,000000 | 0,000000 | 0,00 |
| 262 | ARG | 1081 | GLN | 0,407921 | 0,558416 | 36,89 | 0,415842 | 0,570297 | 37,14 | 0,449505 | 0,335606 | -25,34 |
| 262 | ARG | 1086 | PHE | 0,514851 | 0,362376 | -29,62 | 0,000000 | 0,000000 | 0,00 | 0,481188 | 0,308911 | -35,80 |
| 263 | THR | 1117 | SER | 0,740594 | 0,544554 | -26,47 | 0,544548 | 0,631677 | 16,00 | 0,744554 | 0,547729 | -26,44 |
| 265 | ILE | 1110 | ARG | 0,738614 | 0,762376 | 3,22 | 0,758416 | 0,831683 | 9,66 | 0,716831 | 0,552326 | -22,95 |
| 266 | ALA | 1086 | PHE | 0,600000 | 0,516826 | -13,86 | 0,580198 | 0,512872 | -11,60 | 0,532673 | 0,456622 | -14,28 |
| 266 | ALA | 1110 | ARG | 0,000000 | 0,000000 | 0,00 | 0,000000 | 0,000000 | 0,00 | 0,459406 | 0,382176 | -16,81 |
| 267 | PHE | 1110 | ARG | 0,560396 | 0,613861 | 9,54 | 0,520792 | 0,431683 | -17,11 | 0,477228 | 0,489893 | 2,65 |
| 267 | PHE | 1115 | ILE | 0,944574 | 0,887129 | -6,08 | 0,912871 | 0,899010 | -1,52 | 0,934653 | 0,763404 | -18,32 |

| Hydrogen Bonds | | WT_CYC | ΔF335_CYC | Variation % | WT_VLS | ΔF335_VLS | Variation % | WT_DOX | ΔF335_DOX | Variation % |
|---|---|---|---|---|---|---|---|---|---|---|
| ICH2-NBD2 | life | 497,600 | 349,400 | -29,78 | 303,600 | 376,200 | 23,91 | 501,600 | 461,200 | -8,05 |
| | <N> | 1,367 | 1,307 | -4,36 | 1,151 | 1,451 | 26,14 | 1,309 | 1,217 | -7,09 |
| | DG | -20,078 | -19,129 | -4,72 | -18,746 | -19,387 | 3,42 | -20,198 | -19,201 | -4,93 |

| TMD-NBD interface | | | | WT_CYC | ΔF335_CYC | | WT_VLS | ΔF335_VLS | | WT_DOX | ΔF335_DOX | |
|---|---|---|---|---|---|---|---|---|---|---|---|---|
| ICH3 | | NBD2 | | Frequency | Frequency | Variation % | Frequency | Frequency | Variation % | Frequency | Frequency | Variation % |
| 800 | ASP | 1086 | PHE | 0,734653 | 0,760396 | 3,50 | 0,734653 | 0,720792 | -1,89 | 0,718812 | 0,838371 | 16,63 |
| 800 | ASP | 1087 | TYR | 0,998020 | 0,988119 | -0,99 | 1,000000 | 0,998020 | -0,20 | 0,968317 | 0,988419 | 2,08 |
| 801 | VAL | 1086 | PHE | 0,899010 | 0,914911 | 1,77 | 0,914851 | 0,932673 | 1,95 | 0,891089 | 0,913151 | 2,48 |
| 801 | VAL | 1087 | TYR | 0,489109 | 0,277228 | -43,32 | 0,526731 | 0,453465 | -13,91 | 0,510891 | 0,290996 | -43,04 |
| 802 | SER | 1087 | TYR | 0,685148 | 0,475248 | -30,64 | 0,708911 | 0,575577 | -18,81 | 0,655445 | 0,467719 | -28,64 |
| 805 | ASP | 1044 | TYR | 0,936634 | 0,623762 | -33,40 | 0,922772 | 0,744554 | -19,31 | 0,930693 | 0,534672 | -42,55 |
| 805 | ASP | 1046 | THR | 0,998020 | 0,998020 | 0,00 | 0,996040 | 0,998020 | 0,20 | 0,998020 | 0,994059 | -0,40 |

| Hydrogen Bonds | | WT_CYC | ΔF335_CYC | Variation % | WT_VLS | ΔF335_VLS | Variation % | WT_DOX | ΔF335_DOX | Variation % |
|---|---|---|---|---|---|---|---|---|---|---|
| ICH3-NBD2 | life | 822,600 | 288,000 | -64,99 | 445,000 | 290,000 | -34,83 | 565,400 | 360,200 | -36,29 |
| | <N> | 2,841 | 2,658 | -6,47 | 2,836 | 2,511 | -11,45 | 2,686 | 2,590 | -3,58 |
| | DG | -21,464 | -18,857 | -12,15 | -19,752 | -18,792 | -4,86 | -20,742 | -19,240 | -7,24 |

| TMD-NBD interface | | | | WT_CYC | ΔF335_CYC | | WT_VLS | ΔF335_VLS | | WT_DOX | ΔF335_DOX | |
|---|---|---|---|---|---|---|---|---|---|---|---|---|
| ICH4 | | NBD1 | | Frequency | Frequency | Variation % | Frequency | Frequency | Variation % | Frequency | Frequency | Variation % |
| 905 | ARG | 401 | TYR | 0,217822 | 0,669307 | 207,27 | 0,215842 | 0,641584 | 197,25 | 0,255446 | 0,740949 | 190,06 |
| 905 | ARG | 438 | GLN | 0,215842 | 0,770297 | 256,88 | 0,304951 | 0,661380 | 116,88 | 0,271287 | 0,848459 | 212,75 |
| 905 | ARG | 441 | GLN | 0,964356 | 1,000000 | 3,70 | 1,000000 | 0,998020 | -0,20 | 0,998020 | 0,992266 | -0,58 |
| 908 | VAL | 467 | ARG | 0,568317 | 0,786138 | 38,33 | 0,532674 | 0,790099 | 48,33 | 0,560396 | 0,784506 | 39,99 |
| 909 | SER | 441 | GLN | 0,998020 | 1,000000 | 0,20 | 0,986139 | 0,996040 | 1,00 | 0,980198 | 0,996133 | 1,63 |
| 911 | THR | 467 | ARG | 0,352475 | 0,974257 | 176,40 | 0,398020 | 0,978218 | 145,77 | 0,233664 | 0,974351 | 316,99 |
| 912 | GLN | 464 | ARG | 0,000000 | 0,000000 | 0,00 | 0,576238 | 0,259406 | -54,98 | 0,000000 | 0,000000 | 0,00 |

| Hydrogen Bonds | | WT_CYC | ΔF335_CYC | Variation % | WT_VLS | ΔF335_VLS | Variation % | WT_DOX | ΔF335_DOX | Variation % |
|---|---|---|---|---|---|---|---|---|---|---|
| ICH4-NBD1 | life | 476,000 | 225,000 | -52,73 | 448,200 | 319,000 | -28,83 | 365,200 | 229,800 | -37,08 |
| | <N> | 2,457 | 3,254 | 32,40 | 2,980 | 3,176 | 6,59 | 2,544 | 3,491 | 37,23 |
| | DG | -19,526 | -18,173 | -6,93 | -19,721 | -18,856 | -4,39 | -19,324 | -18,230 | -5,66 |

The variation percentage is estimated based on the mean frequencies of five replica systems for the WT and ΔF335 variant (*holo* systems).
